# The cost and payout of age on germline regeneration and sexual maturation in *Platynereis dumerilii*

**DOI:** 10.1101/2024.01.22.576726

**Authors:** Bria Metzger, B. Duygu Özpolat

## Abstract

Regeneration, regrowing lost and injured body parts, is an ability that generally declines with age or developmental transitions (i.e. metamorphosis, sexual maturation) in many organisms. Regeneration is also energetically a costly process, and trade-offs occur between regeneration and other costly processes such as somatic growth, or sexual reproduction. Here we investigate the interplay of regeneration, reproduction, and age in the segmented worm *Platynereis dumerilii*. *P. dumerilii* can regenerate its whole posterior body axis, along with its reproductive cells, thereby having to carry out the two costly processes (somatic and germ cell regeneration) after injury. We specifically examine how age affects the success of germ cell regeneration and sexual maturation in developmentally young versus old organisms. We hypothesized that developmentally younger individuals (i.e. lower investment state, with gametes in early mitotic stages) will have higher regeneration success and reach sexual maturation faster than the individuals at developmentally older stages (i.e. higher investment state, with gametes in the process of maturation). Surprisingly, older amputated worms grew faster and matured earlier than younger amputees, even though they had to regenerate more segments and recuperate the more costly germ cells which were already starting to undergo gametogenesis. To analyze germ cell regeneration across stages, we used Hybridization Chain Reaction for the germline marker *vasa*. We found that regenerated worms start repopulating new segments with germ cell clusters as early as 14 days post amputation. In addition, *vasa* expression is observed in a wide region of newly-regenerated segments, which appears different from expression patterns during normal growth or regeneration in worms before gonial cluster expansion. Future studies will focus on determining the exact sources of gonial clusters in regeneration.

**Funding:** NIGMS 1R35GM138008-01, Hibbitt Fellowship, WashU Startup funds

## INTRODUCTION

Organisms have limited energy resources at any given developmental stage and therefore need to have trade-offs for where to allocate these resources. Highly regenerative organisms need to balance the costly processes of regeneration, somatic growth, and sexual reproduction (Fields and Levin, 2020; Harshman and Zera, 2007; Henry and Hart, 2005; Partridge et al., 2005). Moreover, these trade-offs may be affected by the developmental stage and age of the organism: older individuals may have different trade-offs than younger individuals due to aging, metabolic, and hormonal state differences. Understanding these trade-offs may shed light on mechanisms that lead to success and failure of regeneration.

Many organisms experience changes in regenerative capacity or reduction and entire loss of this ability as they progress in developmental stages and age (Brunet et al., 2023; Poss, 2010; Sousounis et al., 2014; Yun, 2015). For example, older sponges and corals regenerate less well (Henry and Hart, 2005), fetal mice can regenerate digit tips at faster rates than adult mice (Borgens, 1982; Han et al., 2003; Tower et al., 2022; Yu et al., 2010), and cardiac regeneration ability decreases with age in mice (Haubner et al., 2012; Santos et al., 2020). In *Xenopus laevis* tadpoles, limb and tail regeneration is highly dependent on developmental stage (Patel et al., 2022; Poss, 2010; Yun, 2015). These changes in frogs are thought to be related to hormonal state changes after metamorphosis (reviewed in Yun 2015). In *Ciona intestinalis*, rate of regeneration decreases as the adult body size increases (Dahlberg et al., 2009). While declining regeneration with increasing age seems to indicate younger means more regenerative, being too young may work against regeneration as well: larval stages lack regenerative ability in many animals that can extensively regenerate as adults (Boyd and Seaver, 2023; Henry and Hart, 2005; Jeffery, 2015). Therefore, age and developmental stage are important intrinsic factors affecting regenerative potential in a given species, but which developmental stage or what age is most permissive to regeneration depends on the species.

Injury is generally expected to have an energetic cost and consequently a negative effect on growth and reproduction outputs such as reduced fecundity, lower egg numbers, smaller egg size, or longer time to sexual maturation (Olive and Clark, 1978; Rennolds and Bely, 2023). Regenerating sponges and corals may have reduced fecundity or may remain infertile (Henry and Hart, 2005) due to resource trade-offs between regeneration and sexual reproduction. All together, previous investigations suggest that developmental stage and age generally impairs regeneration in many organisms, and other studies show that injury/regeneration may delay reproductive processes due to resource trade-offs. However, how age, injury/regeneration, and reproduction interact with each other, and whether the trade-off decisions between regeneration and reproduction change in the context of age have been poorly studied.

We use the marine segmented worm (annelid) *Platynereis dumerilii* for investigating the interplay between regeneration, sexual reproduction, and age. *P. dumerilii* can regenerate its posterior body axis upon bisection, allowing us to test the effect of regeneration on reproduction (Fischer and Dorresteijn, 2004; Özpolat et al., 2021; Vervoort and Gazave, 2022). These bisections remove the majority of the germ cells, adding an additional cost to regeneration because the worms have to regenerate both somatic and gametic tissues. Individuals can be staged easily by counting the number of segments, because as most annelids, *P. dumerilii* grows throughout its life by producing new segments at its posterior growth zone located near the tail end.

Therefore, longer worms are developmentally more progressed and aged. Reproductive development and maturation is tightly correlated with the number of segments (reminiscent of pupal weight thresholds in insects required for metamorphosis). For example, worms need to reach about ∼35 segments before they start producing numerous germ cell (gonial) clusters, the earliest stage in gametogenesis (Kuehn et al., 2021; Rebscher, 2014; Rebscher et al., 2007; Zelada González, 2005). As the worms grow longer, gonial clusters start progressing in their gametogenesis, and in worms longer than 60 segments, typically stages of oogenesis in females or spermatogenesis in males are observed (Kuehn et al., 2021; Zelada González, 2005). The worms then need to reach about 70 segments before they can start sexually maturing. Therefore, the biology of *P. dumerilii* enables robust developmental staging in the post-embryonic phase of life. Altogether, developmental staging, the high regenerative ability, and the unique reproductive biology of the species allow testing questions on the interplay of age, regeneration, and reproduction.

Here in this context of the interplay of age, regeneration, and reproduction in *P. dumerilii* we ask how the success of germ cell regeneration and sexual maturation compares between developmentally younger individuals (i.e. lower investment state, with gametes in early mitotic stages) with developmentally older stages (i.e. higher investment state, with gametes in the process of maturation). We hypothesize that older animals that are amputated would be more negatively affected in several ways (measurable by fecundity, time to sexual maturation, regeneration success, and survival) compared to younger amputated animals. We show that while older animals have a higher rate of failure to regenerate, when they do regenerate, contrary to our expectations, they reach sexual maturation significantly faster than younger animals and without any adverse effects. We test whether this is due to higher rate of growth after regeneration, and show that the oldest group has a higher growth rate. We also analyze the gonial cluster regeneration timeline in the youngest and oldest groups using in situ Hybridization Chain Reaction (HCR) using *vasa* mRNA detection as a marker. We show that worms start repopulating regenerated segments with gonial clusters without having to reach the ∼35 segment threshold. We also show that there is a high and broader *vasa* expression that is not typically observed in amputated worms at younger stages. We discuss these results in the context of hormonal regulation that changes with age in *P. dumerilii*, and present some future directions for how to address open questions regarding the hormonal regulation of this process as well as cellular origins of regenerated gonial clusters.

## METHODS

### Github Resources

R code, protocols, and additional resources can be found in a GitHub repository dedicated to this manuscript (ÖzpolatLab-GitHub-Metzger, 2024).

### Animal Culturing

Animals were cultured as previously described (Kuehn et al., 2019) unless otherwise noted. All samples used in this study were kept at light/dark photoperiodic conditions of 16:8h and 8 days of moonlight at night every 28 days. All boxes were checked for deaths and maturations at minimum twice per week. For food, worms received a 1X preparation of spirulina powder and sera micron as follows: 4g spirulina powder and 1.2g sera micron per 2 L of 0.22 µm filtered sea water. The amount of food depended on the density of worms, size of the culturing container, or experiment (specified below in each experiment). For experiments carried out at the Marine Biological Laboratory, natural sea water was collected from the ocean water of Great Harbor and was 1 µm- filtered at the lab for culturing animals (Filtered Natural Sea Water (FNSW)). For experiments in St. Louis, Filtered Artificial Sea Water (FASW) was prepared with Red Sea Salt (RSS, Red Sea Fish) to a salinity of 35 ppt. Prior tests by our lab and other *Platynereis* labs established the suitability of RSS sea water and found no significant difference in growth rate or time to maturation.

#### Amputation & fecundity (A&F) experiment (batch culturing)

Animals were kept in plastic sterilite culture boxes with 500mL of FNSW. At the beginning of the experiment, each experimental and control box had between 13 and 18 animals. All boxes received a complete water change once every two weeks, and death and mature checks were performed at least three times per week. The number of worms per box was checked and updated at least weekly, with additional updates each time a worm was discovered dead or mature and removed. The amount of food was scaled to the current number of worms in the box such that each worm received 333 µL of 1X spirulina and sera micron mix per feeding.

#### Single worm growth rate experiment (individual culturing)

Animals were kept individually in 6-well plates with 7 mL of 0.22 µm FASW and final concentration of 1,000 U/mL penicillin and streptomycin (Fisher Scientific, 15140122). Feeding began 3 days post amputation to allow for the gut to reform; unamputated controls also began feeding 3 days after the beginning of the experiment. Each worm received 333 µL of 1X spirulina and sera micron prepared food on the regular feeding schedule. Once per week, animals received a complete water change and were moved to fresh wells. To protect against evaporation, well plates were kept in a humid chamber: a large plastic box with a loose lid and a damp paper towel lining the bottom. To prevent mold growth, the paper towel was changed out weekly alongside water changes.

### Amputations

We amputated worms within three ranges: 40-49 segments, 50-59 segments, and 60-69 segments. In later sections, these range groups are referred to as 40s, 50s, and 60s. These ranges were selected to encompass the range between germline expansion (40s) and just prior to the onset of sexual maturation (60s). Segments were counted according to ‘Segment Counts’ below.

For all amputations, worms were anesthetized for 15 minutes in 1:1 7.5% MgCl2:0.22 µm FNSW or FASW, then rinsed briefly in sea water before being transferred to a glass slide. In most experiments amputations were performed between the 8th and 9th chaetigerous segment. For the experiment testing the effect of amputation location, the amputations were performed between the 7th and 8th segment, between the 8th and 9th segment, or roughly 2/3rds down the body of the worm. For the experiment assessing fecundity after amputation, any worms that died before posterior regeneration was completed (5 DPA) were discarded and noted, but not factored into the long-term total mortality counts of the experiment. For experiments involving single worm cultures, mortalities from the start of the experiment (0 DPA) were factored into total mortality.

#### A&F experiment

Most amputations were performed over the course of three days: all 40-segment amputations on the first day, followed by all 50-segment amputations on the second and all 60-segment amputations on the third to minimize any effect of uncontrollable fluctuations in culture conditions on recovery. In the case of 60-segment amputations, one fertilization group showed fungal contamination and mass mortality. All individuals from this fertilization group were discarded and a fresh series of 60-segment amputations was performed 4 weeks later and incorporated into the experiment. To recover, the amputees were kept individually in 6-well plates with 7mL of 0.22 µm FNSW/FASW and a final concentration of 1,000 U/mL penicillin and streptomycin. After 6 days in 6-well plates, the worms were transferred into 750 mL culture boxes and cultured as described above.

#### Single worm segment addition rate experiment

All 40-segment worms were amputated on the first day, followed by all 60-segment worms the day after to minimize any effect of uncontrollable fluctuations in culture conditions on recovery. 40- and 60-segment group amputations were performed in triplicate with each set of amputations occurring ∼1 month apart. Animals were kept individually in 6-well plates for the duration of the experiment and cultured as described above. Worms were staged and segment-counted weekly, which allowed us to track individual trajectories of posterior regeneration, growth, and mortality. Segment counts were tracked until 63-64 days (9 weeks) after amputation. We chose this time frame based off of the segment counts observed in our previous experiment (Suppl. Fig. 2), which suggested a rapid period of growth around 28 DPA and a slower period of growth by 63 DPA. We expected this time frame to be most useful for revealing age-specific differences in segment addition that could impact time to maturation.

### Segment Counts

Worms were anesthetized for 15 minutes in 1:1 7.5% MgCl2:0.22 µm FNSW or FASW, rinsed briefly in sea water, then imaged using a Zeiss Stemi 305 CAM BODY stereoscope with a built-in camera and an iPad with Lab Scope. These images were used to conduct segment counts using the Cell Counter plugin in FIJI. We counted each chaete-bearing segment, excluding segment primordia that had not yet formed chaete.

In rare cases, worms auto-amputated close to the beginning of the experiment. If a worm had a visible amputation plane or a reduction in segments counted from one time point to the next, it was removed from analysis. One 40 segment control worm and three 60 segment control worms that auto-amputated were removed from analysis.

All segment counts are provided as csv files (see Suppl. File 1 for description of files, Suppl. File 2 for R codes, and Suppl. File 3 for the csv files).

### A&F Experiment Analyses

#### Assessment of Maturation

For each mature individual, we recorded the date of maturation, the total number of segments at maturation, and the location of the atokous/epitokous boundary, determined by the last segment displaying atokous chaetae (Schulz et al., 1989). Mature animals were anesthetized and imaged as described above.

#### Assessment of Fertility and Fecundity

After imaging, mature animals were paired for fertilizations. When possible, mature animals from the experiment were paired with mature animals from our stock cultures. This pairing allowed us to separate any sex-specific effects on fertility and fecundity, assuming a stable rate of fertility in the stock cultures. For example, if all fertilizations between amputated females and stock males failed but all fertilizations between amputated males and stock females succeeded, we would conclude that amputation impairs female, but not male, fertility. Fertilizations between amputated animals would obscure potential sex-specific differences. In one instance, no mature animals were available from the stock culture and one (successful) fertilization was staged between two 50-segment amputated mature animals.

Infrequently, a female spawned but a male failed to release sperm naturally; in these cases, the male was prodded with a pipette to release sperm, and failing that, cut open with a scalpel on a glass slide so that the contents could be rinsed into the fertilization dish to assist fertilization. Males from stock cultures as well as amputated and control groups occasionally failed to spawn without assistance, so failure to spawn naturally does not necessarily indicate infertile or defective individuals, as spawning behavior itself is ruled by complex mating cues that are yet to be fully understood.

1-4 hours after fertilization, the eggs were collected and de-jellied in an 85 μm mesh filter and rinsed a minimum of 10 times with 0.22 μm FNSW, with additional rinses if jelly remained. Then, we prepared the eggs for quantification and assessment of fecundity. The eggs were resuspended in 50 mL of 0.22 μm FNSW in a 50mL plastic beaker, then poured rapidly between two 50 mL beakers to achieve an even suspension. A 1 mL aliquot was removed from this suspension and dispensed into a watch glass. The eggs were viewed under magnification and counted using a watch glass. Fertilized eggs were distinguished from unfertilized eggs based on their color: unfertilized eggs became yellowed, clouded, and finally opaque, while fertilized eggs became clear and began cleaving. The total number of eggs, the number of fertilized eggs, and the number of unfertilized eggs were recorded for each individual assessed.

All scores and counts are provided as csv files (see Suppl. File 1 for description of files, Suppl. File 2 for R codes, and Suppl. File 3 for the csv files).

### Fixations

Worms were anesthetized as described above and briefly rinsed in FNSW/FASW. Then, they were submerged in fixative pre-chilled to 4℃. The fixative was composed of 1 part 16% paraformaldehyde, 2 parts 2X PBS made with DEPC-treated water, and 1 part 0.22 µm FSNW/FASW. The worms were fixed on an orbital shaker for 2 hours on ice or overnight at 4℃. Following fixation, worms were dehydrated through 15-minute washes of 25%, 50%, 75%, and 100% methanol in DEPC-treated PBSt, chilled to 4℃. Worms were stored in fresh 100% methanol at -20℃.

### Hybridization Chain Reaction (HCR)

#### Probe design

Probes were designed through the Probe Maker software we developed (Kuehn et al., 2021). The sequences generated by the software were used to order a single, batched DNA oligo pool (50 pmol DNA oPools Oligo Pool) from Integrated DNA Technologies, resuspended to 1 pmol/μl in 50 μL nuclease-free water (Ambion, 10977015). B3 hairpins conjugated to Alexa-647 were purchased from Molecular Instruments. Probe sequences are provided in the supplementary information (Suppl.

File 04).

#### Tissue collection and HCR

To visualize gonial cluster regeneration in worms of different ages, we amputated worms between 40-49 (40s) and 60-69 (60s) segments and allowed them to regenerate until 7 DPA, 14 DPA, or 23 DPA. At each of these timepoints, a subset of worms were fixed, dehydrated in a methanol series, and stored in 100% methanol at -20℃.

Probeset for *vasa* were designed using the HCR Probe Maker (Kuehn et al., 2021). Probe hybridization buffer, probe wash buffer, amplification buffer were sourced from Molecular Instruments (https://www.molecularinstruments.com) (Choi et al., 2014). HCR 3.0 was performed according to the published methodology (Choi et al., 2018) with a few modifications to suit our samples as previously described (Kuehn et al., 2021). Additionally, in place of proteinase K digestion followed by post-fixation, permeabilization was achieved through a 30 min wash in a Tween-based detergent solution (Bruce et al., 2021). At the amplification step, DAPI was added to a final concentration of 2 μg/ml. The full protocol adapted for *Platynereis* is provided in the GitHub repository (ÖzpolatLab-GitHub-Metzger, 2024).

#### Imaging

HCR samples were mounted in Slowfade Glass with DAPI (Thermo-Fischer S36920-5X2ML), kept at 4°C until imaging, and imaged using Zeiss Laser Scanning Confocal Microscope LSM 900 using Z-stacks and tiling. For each experiment, a no-probe, hairpin-only control was included to support differentiation of true signal from baseline background and autofluorescence. To survey all possible gonial clusters in original and regenerated segments, we acquired tiled Z stacks encompassing the region immediately posterior to the jaw to the pygidium. A step size of 8 µm was chosen so that all germline structures would be captured in a minimum of one slice. All stages of female germline structures exceed 8 µm (Fischer, 1975). Male germline structures exceed 8 µm up through the spermatocyte stage, which average 6.7 µm in diameter (Meisel, 1990), and no male worms were observed past the spermatogonia ball I stage.

#### Image analysis

Images were processed for levels and annotated with scale bars in FIJI (Schindelin et al., 2012) or Imaris (Bitplane). Panels were assembled in Adobe Illustrator. For segment counts, only chaetae-bearing segments were counted (new segments without chaetae were not counted). For gonial cluster counts, if the anterior cluster was the dispersed phenotype, these were counted as gonial clusters, otherwise, if the anterior cluster was the ring or stripe phenotype, these were not counted as gonial clusters. Single oocytes were included in gonial cluster count. However, cells within the broad *vasa* expression, or cells that do not have gonial cluster or gamete phenotype were not included in the count. For regeneration stages, previously-established staging for *P. dumerilii* was used (Planques et al., 2018). Scoring table of results is provided as supplementary material (Suppl. File 5)

### Phalloidin Staining

28 DPA, a 60s-amputated worm from the segment addition rate experiment was observed to be dying, signified by minimal response to stimuli, tissue deterioration, and depigmentation. To explore potential factors in 60s mortality, this worm was removed from the experiment and placed into fixative (4% PFA in ASW-PBSt, 2 weeks nutating at 4C). The worm was then stained with DAPI (1 ug/100 µL) and Phalloidin (Alexa-647, 1X) together in PBS for 45 mins at room temperature. To aid in mounting, the head and pharynx, including the first four segments, were removed. The tail segments were mounted and imaged as above (‘Imaging’).

### Plots and Statistical Analyses

Line graphs, violin plots, and bubble plots were generated in RStudio (R Development Core Team, 2011; RStudio Team, 2020) using ggplot 2 (v3.3.3) (Wickham, 2016). Visuals were collected and edited in Adobe Illustrator for clarity, shape, axis labels, and text size adjustment. ANOVAs, paired t tests, and Wilcoxon non-parametric tests were performed in RStudio.

For the time to maturation and number of segments at maturation, univariate and multivariate ANOVAs were performed by the Biostatistics Consulting Service at Washington University in St Louis. The data analysis was generated using SAS software, version 9.4 of the SAS System for Windows (SAS Institute Inc., Cary, NC, USA). In univariate models of segment group, amputation, and sex, differences among segment groups were highly significant (p<0.0001), differences between amputated and control groups were marginally significant (p=0.0857), and males and females were not significantly different (0.2857) We then ran a multivariate model including predictors in univariate models with p<0.1, which included segment group, amputation, and their interaction. In the multivariate model, the interaction between segment group and amputation was marginally significant (p=0.0584). We performed all planned comparisons. P-values for multiple comparisons were adjusted with the Tukey-Kramer adjustment.

All R scripts and datasets are available as csv files (Suppl. File 1 for description of files, Suppl. File 2 for R codes, and Suppl. File 3 for the csv files) and also via GitHub (ÖzpolatLab-GitHub-Metzger, 2024).

## RESULTS

### Amputated *Platynereis dumerilii* reach sexual maturation, are fertile, and show normal reproductive morphology following regeneration

*Platynereis dumerilii* can regenerate the whole posterior body axis when bisected, but they fail to regenerate anteriorly. Worms also fail to regenerate if the head is smaller than 8 chaetigerous segments (Suppl. Fig. 1). Amputations leaving 8-segment or longer head pieces will typically result in body axis regeneration. As the germ cells are stored along the body cavity, such amputations also remove the majority of the germ cells, which regenerate along with body axis regeneration (Fig. 1A-B). We wanted to test if worms that had to regenerate their germline were impacted on their sexual maturation, morphology, and reproductive success. To characterize the success of sexual reproduction following amputation and regeneration, we compared the maturation of amputees and controls of different initial segment lengths that span the period between germline expansion (i.e. when germ cell clusters start appearing in the coelomic cavity) at ∼35 segments (Kuehn et al., 2021) and sexual metamorphosis at ∼70 segments (Fig. 1A). We split this period into three segment ranges to capture approximate phases in germline development: 40-49s (the stage when small gonial clusters are becoming numerous), 50-59s (when gonial clusters are generally in the early stages of gametogenesis), and 60-69s (when the gonial clusters are generally in late stages of gametogenesis, or full maturation is imminent) (Fig. 1B). For each individual, the amputation was performed after the 8th pair of chaetae, the most anterior possible cut site for which regeneration proceeds without excessive mortality in previous works (Planques et al., 2018). Our pilot experiments on worms shorter than 50 segments showed that, by 7 DPA, worms amputated after the 8th segment regenerated comparably to worms amputated after the ∼30th segment, while worms amputated after the 7th segment were significantly delayed or unable to close the wound (Suppl. Fig. 1A). Therefore we proceeded with amputations after the 8th segment to maximize the removal of segments containing the germ cells. However, it did not remove the anterior germline cluster, which is typically located between the 5th and 6th segment. Amputated worms were allowed to regenerate, and both amputated worms and their controls were raised to sexual maturation (see Methods for details).

**Figure 1:**
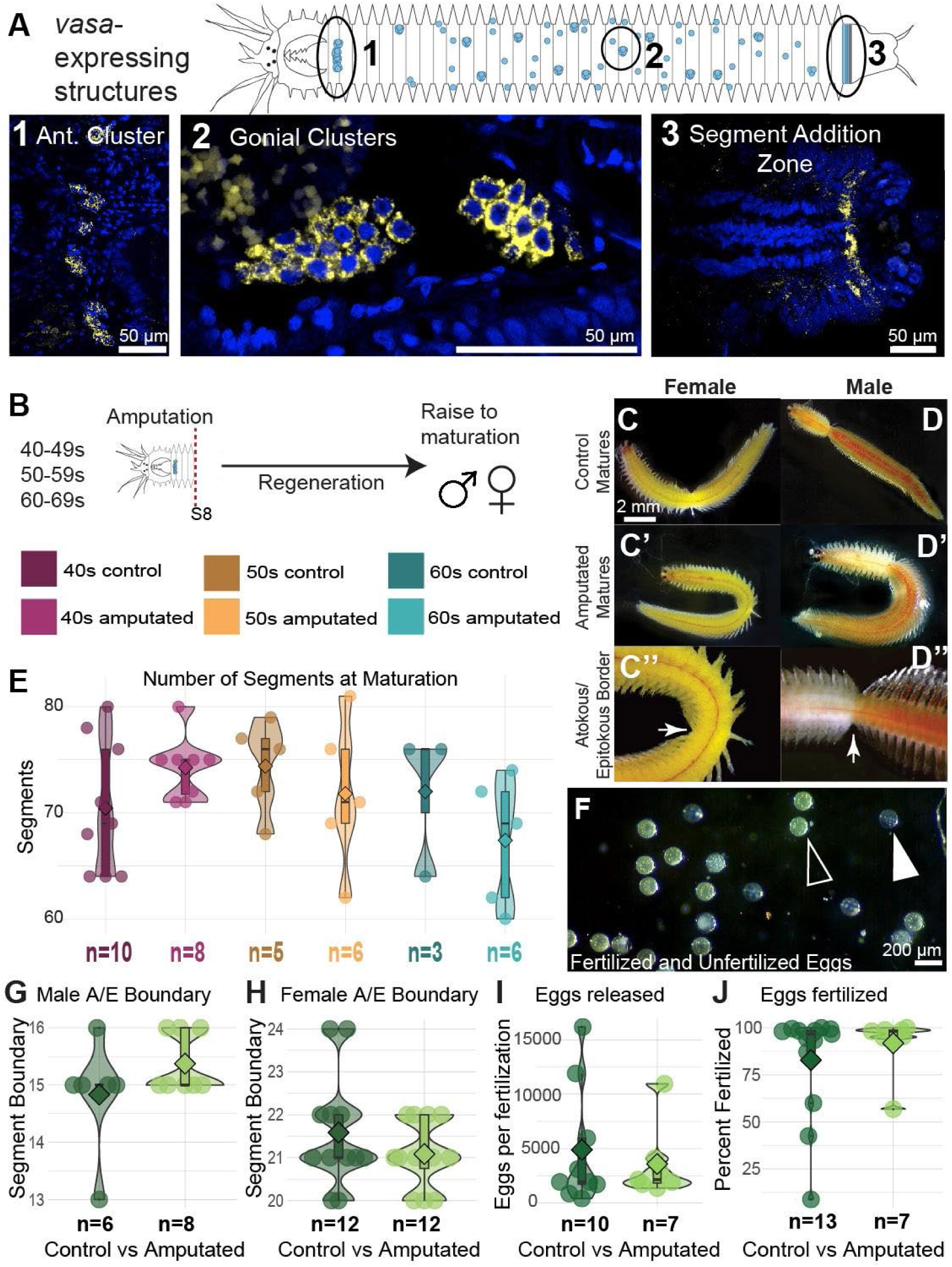
Amputated worms have morphologically and functionally normal maturation after germline regeneration. A) Example worm showing the *vasa* expressing structures (yellow). An anteriorly-located band of germ cells (1) is thought to be the source of the gonial clusters (2) which spread across the worm’s body and float in the coelomic fluid. DAPI stain in blue. Scale bars 50 μm. B) Experimental schematic. Worms were collected into three groups of 10-segment ranges: 40-49 segments, 50-59 segments, and 60-69 segments. These ranges span the developmental period between germline expansion (∼35 segments) and sexual metamorphosis (∼70 segments). Worms were amputated after the 8th pair of chaetae and raised to maturation alongside unamputated controls. C) Control mature female. Scale bar is 2 mm for C-D’. C’) Amputated and regenerated mature female. C’’) Female atokous-epitokous border, indicated by white arrow. D) Control mature male. D’) Amputated and regenerated mature male. D’’) Male atokous-epitokous border, indicated by white arrow. E) Number of segments at maturation. In univariate models, there was no difference among the 3 segment group (p=0.3115), between the control and amputated groups (p=0.9127) or between males and females (p=0.2257). F) Example photo showing how fertilized (solid arrowhead) and unfertilized eggs (outlined arrowheads) can be visually differentiated by color and opacity by 2 hours post fertilization. Scale bar 200 μm. G-H) The location of the male and female atokous/epitokous border did not differ significantly between control and amputated matures (Wilcoxon signed rank test on paired samples, p=0.285 for males, p=0.464 for females). I) The number of eggs released upon spawning did not differ significantly between control and amputated mature females (Wilcoxon, p=0.812). J) Percentage of eggs fertilized did not differ significantly between control and amputated matures (Wilcoxon, p=0.751).

**Figure 2:**
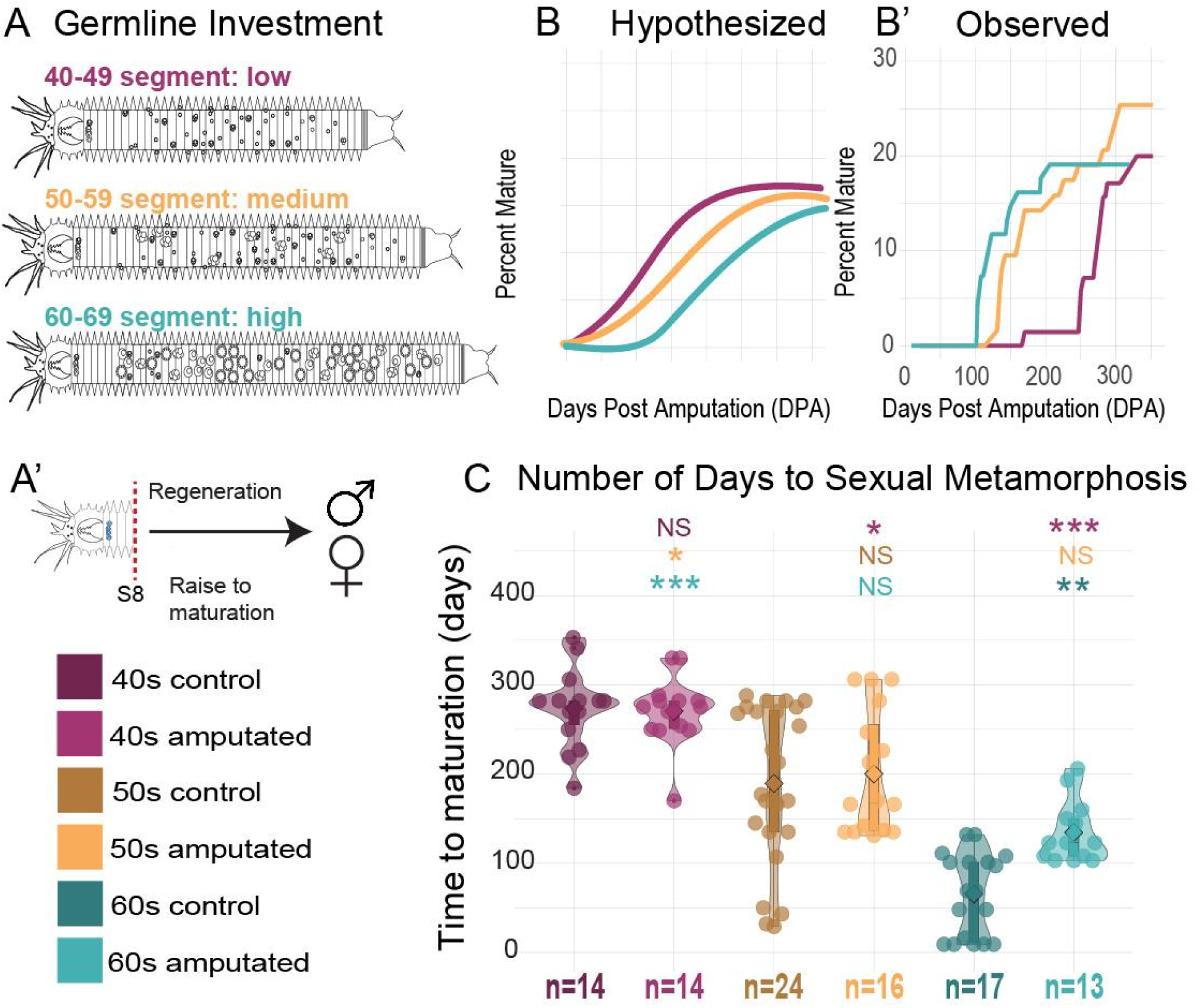
Developmentally “older” regenerates mature earlier than “younger” regenerates. A-A’) Experimental schematic. Worms of three 10-segment ranges spanning low, medium, and high germline investment (A) were amputated after the 8th pair of chaetae and raised to maturation alongside unamputated controls (A’). B-B’) Hypothetical vs observed results. We hypothesized that older worms would take longer to maturation, however we observed the opposite result. C) Violin plots showing the number of days to reach maturation per group (controls and amputated). 60s amputees matured the earliest at an average of 134 days to reach sexual metamorphosis, followed by 50s amputees at an average of 200 days, and 40s amputees at an average of 270 days. Time to maturation differed significantly between 60s amputees and 40s amputees (p<0.001). While 50s amputees did not differ significantly from 40s or 60s amputees, this group was in between the other two groups, showing a gradual shift. Asterisks indicate the level of significance of planned pairwise comparisons (* p<0.05, ** p<0.01, *** p<0.001, NS if p>=0.05).

Overall, amputated worms did not show any significant differences from control worms in fecundity, fertility, and reproductive morphology (Fig. 1C-J). We assessed the production of functional gametes after regeneration, by scoring for the number of eggs released and the percentage of eggs fertilized (Fig. 1F). Mature animals from within the experiment were paired for fertilization with matures from the wild type cultures when possible. There were no significant differences in the number of eggs released or the percentage of eggs fertilized (Fig 1I-J; Wilcoxon signed rank test, p =0.8596 for eggs released, p =0.4787 for eggs fertilized). From these fertilizations, F1 and F2 offspring were raised to maturation and reproduced successfully as well (data not shown).

Previous literature indicates that *Platynereis dumerilii* typically undergoes sexual metamorphosis between 65-75 segments (Fischer, 1975, 1974; Legras et al., 2023). However, whether the number of segments at maturation differ in amputated-regenerated worms have not been analyzed systematically. It is possible that amputated worms would reach a similar segment threshold for maturation, or alternatively, amputation and subsequent regeneration would alter or decouple the relationship between segment number and transition to sexual maturation. We compared a subset of our control and amputated groups by counting total segment number at maturation. In both control and amputated worms we observed a more flexible segment range at maturation than previously-reported results, with worms being between 60 and 82 segments at maturation (Fig. 1E). Univariate models showed no significant difference in the number of segments at maturation among different segment groups (p=0.3115), between the control and amputated groups (p=0.9127) or between males and females (p=0.2257; Suppl. Fig. 1B-C). These data indicate that regenerated worms reach a similar number of segments for sexual metamorphosis as control worms.

We also assessed whether the segment-specific location of a mature characteristic, the atokous-epitokous border, would be maintained in mature regenerated animals. In *P. dumerilii*, sexual metamorphosis involves dramatic restructuring of the somatic tissues. A sex-specific number of parapodia at the anterior end of the worm maintain an “atokous” phenotype, while all parapodia posterior of this boundary take on an “epitokous” phenotype with transformed muscles and knife-like chaetae. This boundary is easily distinguishable in males by the boundary between the white, sperm-rich anterior and the red muscular posterior (Fig. 1D’’). In females, morphology of the parapodia is sufficient to identify the border (Fig. 1C’’). In males, the atokous-epitokous boundary occurs after the 14th-16th segments, while in females, the atokous-epitokous boundary occurs between the 20th-24th segments (Schulz et al., 1989). For all amputated groups, we found that the atokous-epitokous border was within the correct sex-specific segment range (Fig. 1G, 1H). For the control groups, one 40s male had the atokous-epitokous border one segment anterior to the normal location, after the 13th segment (Fig. 1G). All others were within the sex-specific segment range.

### Amputated worms show stage-specific timelines of regeneration and maturation

We next wanted to analyze if stage of development (measured by segment number) had an effect on time to maturation after amputation. We expected that worms of different developmental stages (40-49 segments, 50-59 segments, and 60-69 segments) would vary in the time required to regenerate all lost somatic and germ cells and reach sexual metamorphosis (Fig. 2A-A’). For brevity, we will refer to these groups as 40s, 50s, and 60s groups. Our rationale was that overall, older worms (e.g. 60s) would have invested more resources into germ cells such as having vitellogenic oocytes, compared to younger worms (e.g. 40s). Therefore, these older worms had to recover a more “expensive” loss, and a longer body axis upon amputation. We expected a gradually-increasing delay to maturation in older worms compared to younger worms (Fig. 2B).

Strikingly however, our analyses revealed the opposite: we found that the older amputated worms matured earlier than younger control and amputated worms, and this shift is gradual as indicated by the 50s amputees maturing sometime between 40s and 60s amputees (Fig. 2B’, C). We ran univariate and multivariate analysis of variance to determine if the days to maturation were different between the 2 treatments (amputated vs control), 3 segment groups (40s, 50s or 60s) and sex at maturation (Male vs Female). In univariate models, there were overall significant differences among the 40s, 50s, and 60s groups (p<0.0001). The treatment groups (amputated vs control) were marginally different (p=0.0857), and there was no significant difference between males and females (p=0.2857). With segment group and amputation as our two independent variables, we ran a multivariate model with an additional interaction term. This allowed us to assess effects of segment group and amputation alone, as well as effects that emerge from combinations of these variables. In the multivariate model, the interaction between segment group and treatment was marginally significant (p=0.0584), meaning that amputation had different effects on different segment groups. Taken together, these data indicate an inverse relationship between the age/developmental stage at amputation and the length of time to regenerate and reach sexual metamorphosis.

Interestingly, when amputated groups were compared with their unamputated controls, we found that for 40s, there was no significant difference in the number of days to reach maturation between the amputated group and the controls (p=1.0) (Fig. 2C). The same was observed for 50s amputees vs controls (no significant difference, p=0.9941), even though the time to maturation showed greater variance for 50s amputees than for 50s controls (Fig. 2C). Effectively, having an amputation that removed the major percentage of the body, and having had to regenerate these missing segments cause no overall delay to maturation in the 40s and 50s amputees compared with their controls. In contrast, 60s amputees were significantly delayed to maturation relative to unamputated controls (p=0.0309), with amputees maturing at an average of 134 days and controls maturing at an average of 65 days (Fig. 2C). As noted earlier, however, even though 60s amputees matured later than their 60s controls, they still matured earlier than both 40s and 50s controls, despite needing to regenerate 52 segments along with their germ cells.

### Segment addition rates differ between 40s and 60s amputees

Since the 60s amputees were maturing earlier, one explanation could be that worms in this group start maturing without growing to 70-80 segments, and instead they can mature at shorter segment numbers compared to the 40s and 50s groups (i.e. adding segments at a comparable rate and just reaching fewer segments before maturing).

However, all groups reached a similar segment number threshold before maturing (Fig. 1E, p=0.07946), and there was not a significant correlation between the length at maturation and the time to maturation (Suppl. Fig. 1D). Another explanation for the 60s amputated group reaching maturation faster could be faster rate of regeneration and segment addition after amputation, allowing the worms to grow to sexual maturation faster. For this experiment, we had collected data on the number of segments at different checkpoints (28, 63, 96 DPA) between amputation and maturation in these batch cultures (Suppl. Fig. 2C-D). The number of segments in amputated groups differed as early as 28 DPA, at which point 40s amputees were significantly shorter than both 50- and 60 segment amputees (p<0.001, p<0.001;). By 84 DPA, 40s amputees were still significantly shorter than 50s and 60s amputees (p<0.001) as well as their unamputated controls (p<0.001), while 50- and 60s amputees had caught up to their controls (p=0.06031, p=0.1909). These data suggest that differential rates of segment addition, established at least as early as 28 DPA, could at least partly underlie the observed differences in the time to maturation. However, because these worms were kept in collective culture, the growth rates of individual worms could not be assessed. Furthermore, collective culture increases the incidence of tail breakage and territorial encounters (and potentially cannibalism, or differential access to food resources due to social and territorial behaviors).

To get a clearer understanding of stage-specific differences in the rate of segment addition following amputation, we performed a follow-up experiment with single-cultured worms, and observed their growth individually across time. We amputated 40s and 60s worms and kept them in individual wells, counted their segments at several time points to assess the rate of growth, and noted mortality (see Methods for details). The growth trajectories of single worms reveal that by 63 DPA, controls grow at a much slower rate than amputees (Fig. 3A-B), and 60s amputees added more segments than 40s amputees (Fig. 3B, F, Tukey-Kramer test, p adj <0.05). Notably, the rate of segment addition is not constant throughout the 63 day time period. Both amputated groups show an earlier, faster period of growth around 20 DPA as well as a later, slower period of growth by 63 DPA (Fig. 3B, D). The period of highest segment addition for both amputated groups appears to occur until ∼30 DPA (Supp. Fig. 3). Until this point, 60s amputees added segments at a higher rate than 40s amputees. After ∼30 DPA, segment addition rates declined and appeared to converge.

**Figure 3:**
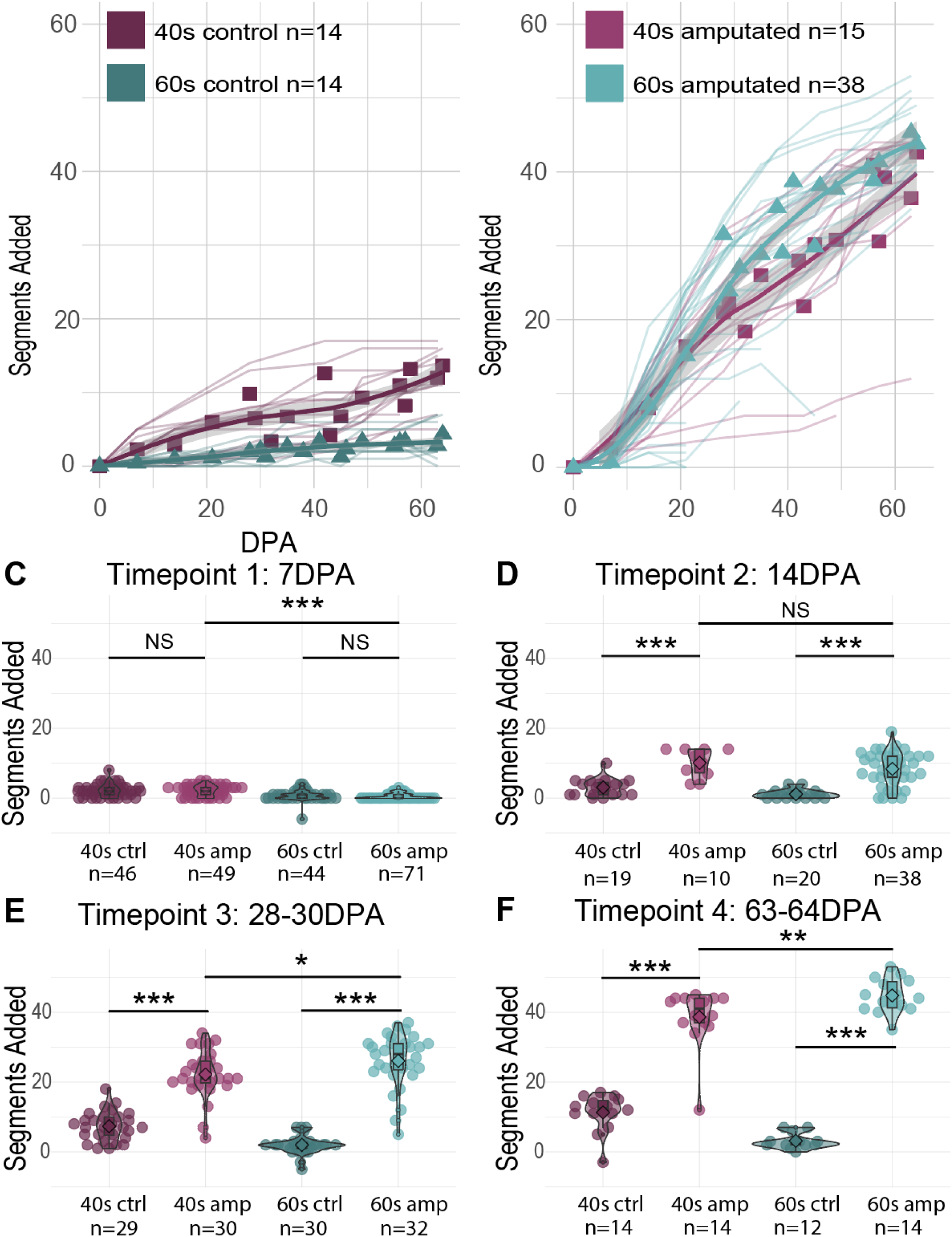
Older (60S) and younger (40S) worms differ in long-term regenerative growth trajectories. A-B) Line graphs show the number of segments added since the beginning of the experiment in days post amputation (DPA) for control and amputated groups. Individual growth trajectories are depicted as thin semi-transparent lines. Triangles represent the means at each point for which data is available (days when segments were counted). A locally weighted regression line is drawn in bold with a 90% confidence region in transparent gray. (Note that graphs in A-B show a detailed subset of data from plots in C-F. In A-B, the segments of individual worms were counted up to 64 DPA. Plots in C-F include segment counts from similarly treated worms at different timepoints (however not all these samples were raised to 64 DPA therefore numbers of samples change)). C-F) Violin plots show the distribution of the number of segments added per group at 4 time points spanning the duration of the experiment, revealing changes in the pace of segment addition among and between groups. Individuals are represented by semi-transparent dots, and violins with box plots superimposed illustrate the distribution within each group. A diamond shows the group mean. A multivariate ANOVA was performed assessing the impact of segment group (40-49 segments or 60-69 segments), amputation (amputated or control), and interaction between these terms. T tests were performed with Benjamini-Hochberg correction for multiple comparisons. Asterisks indicate the level of significance of pairwise comparisons (t tests; * p<0.05, ** p<0.01, *** p<0.001, NS if p>=0.05). At 7 DPA, 40s amputees have added more segments than 60s amputees (p=2.40E-06). By 14 DPA, amputees no longer significantly differ from each other (p=0.20). By 28 DPA, 60s amputees have added more segments than 40s amputees (p=0.01058), which persists as growth appears to plateau approaching 63 DPA (p=0.0072). One 40s control sample and three 60s control samples auto-amputated 29 days after the beginning of the experiment and were removed from analysis.

Interestingly, a finer-scale analysis of segment numbers shows that 60s amputees did not immediately begin adding more segments than 40s amputees. In contrast, they were significantly delayed early on (Fig. 3C). At 7 DPA, 40s amputees have added significantly more segments than 60s amputees; 40s amputees have added an average of 2.6 segments compared to an average of 0.55 segments for 60s amputees (Fig. 3C, p=2.4e-06). Proportionally, 93% of 40s amputees have added at least one segment, indicating successful regeneration of the posterior growth zone by 7 DPA, compared to only 39% of 60s amputees. Previous work has established that the normal timeline for regeneration of the posterior growth zone occurs within 5 days, culminating in late patterning/visible segment formation. Images of 7 DPA regenerates show that 40s amputees are reaching the expected stage of regeneration by 7 DPA, but 60s amputees show delayed initiation of segment addition as well as high mortality (see below). Despite this initial delay, by 14 DPA, surviving 60s amputees caught up to 40s amputees and the differences between amputated groups were no longer significant (Fig. 3D, p=0.20). By 28 DPA, 60s amputees have added more segments than 40s amputees (Fig. 3E, p=0.01058) and continue to outpace 40s amputees until approximately 35 DPA, at which point the rates appear to converge (Suppl. Fig. 3). Overall, 60s amputees generally show a faster segment addition rate compared with the 40s controls, which could partly explain the faster timeline of getting to sexual maturation in this group.

We expected amputated worms to have an elevated segment addition rate compared to controls, given that regenerating worms have been shown to add segments faster than unamputated controls (Gazave et al., 2013). Our data exemplify this in both age groups: by 63 DPA, 40s amputees have added an average of 37.0 segments in comparison to an average of 11.0 segments in 40s controls (Fig. 3F). The difference between 60s amputees and controls is even greater, with 60s amputees adding an average of 44.2 segments and 60s controls adding an average of 3.2 segments over the experimental time course (Fig 3F). By the end of the 63 DPA, segment addition rates of amputated worms are declining, but still slightly elevated compared to controls (Supp. Fig. 3).

### Amputated worms show age-specific mortality, regeneration defects and delays

While the surviving 60s amputees grow and mature faster, this group also had a higher rate of mortality and regeneration defects. The mortality rate of 60s amputees was highly elevated compared to 60s controls and 40s amputees. We carried out comparisons in the single-worm culture experiment and found that over the 63 day course of the experiment, 57.5% of 60s amputees died compared to 16.67% of 60s controls (Fig. 4A). The mortality rate for 40s amputees was only 6.25%, and no 40s controls died (Fig. 4A). Instances of death were elevated earlier within 7 DPA, but otherwise distributed across the experimental timeline. 8 of 23 total deaths (34.8%) among 60s amputees occurred on or before 7 DPA (Fig. 4B). This early elevated mortality occurred alongside delays and failures of posterior regeneration, and seemed to coincide with or were possibly precipitated by the first feeding after amputation.

**Figure 4:**
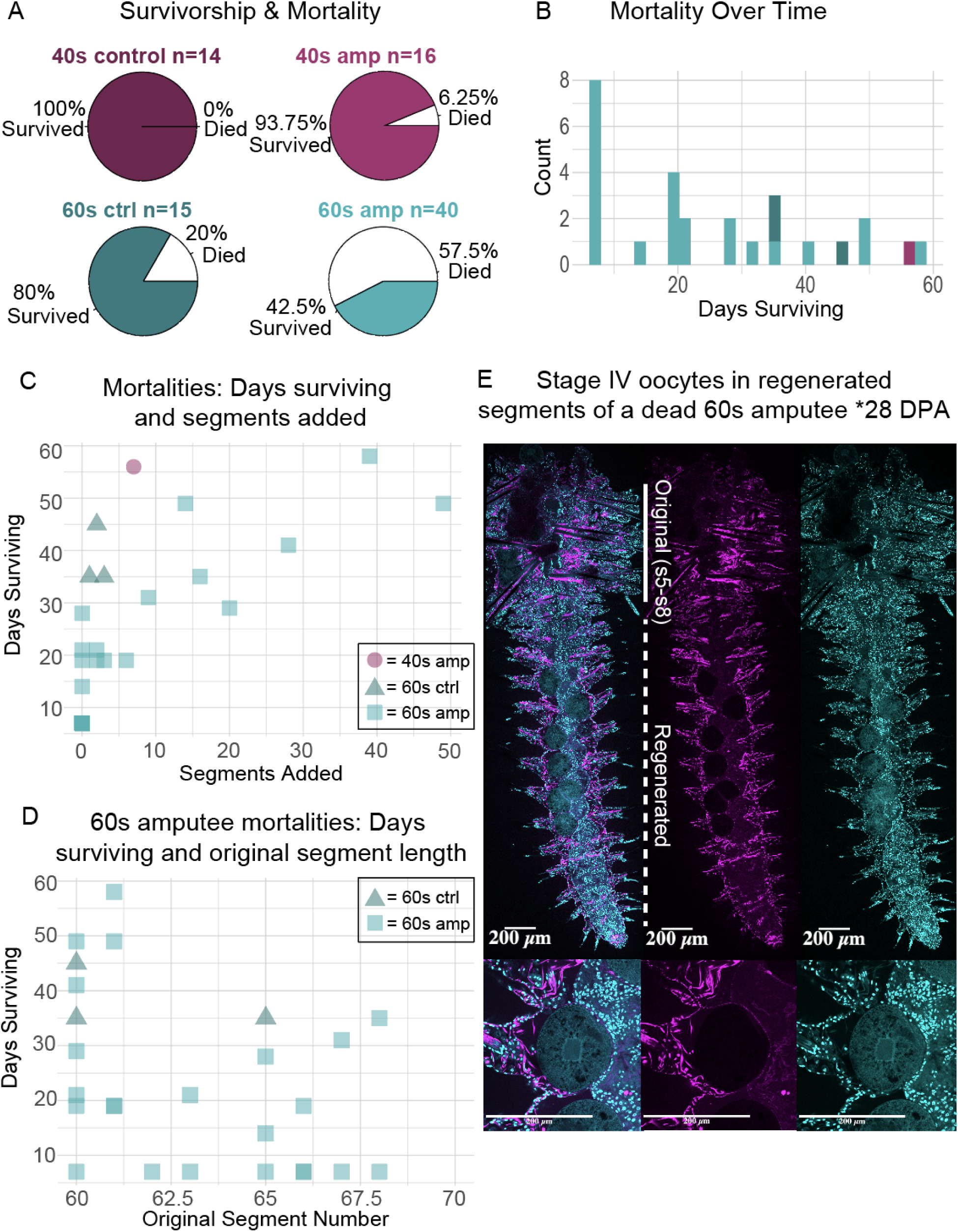
Age-specific mortality, regeneration defects and delays. A) Pie charts showing the percentage of survivorship and mortality for each group at the end of 63 DPA. Survivorship is represented by color and mortality is represented by white space. B) Stacked histogram of mortality in each group over time. 60s amputees have elevated mortality compared to all other groups. C) A scatterplot illustrating the number of days surviving and the original number of segments for all worms that died 0-64 DPA. D) A scatterplot illustrating the number of days surviving and the number of segments added for all 60s worms that died 0-64 DPA. E) A fixed 60s amputee stained with DAPI (cyan) and phalloidin (magenta) to illustrate the presence of mature oocytes in the regenerated segments. This individual survived until 28 DPA, had regenerated successfully, and had added 16 segments by the time of death. Original segments are denoted by the solid white bar, and regenerated segments by a dashed white bar. All scale bars are 200 μm.

Most 60s amputees that died after 7 DPA had added several segments, indicating functional regeneration of the growth zone and gut structures (Fig. 4C). However, there were also several 60s amputees that survived for weeks without adding segments (Fig. 4C). Among 60s amputees, there does not appear to be a correlation between original segment number and the number of days surviving (Fig. 4D), suggesting that there is not a sharp threshold for survival following amputation, or progressively early mortality with increasing segment number. It is unclear why these worms, which had successfully added several segments, died. We noted that in one of these dying 60s amputees there were nearly mature oocytes in the regenerated region (Fig. 4E). Previous studies indicate that submature oocytes may exert negative feedback on growth and regeneration (Hofmann, 1975; Porchet and Cardon, 1976). Notably, many 60s amputees that successfully regenerated and survived for the full 63 days were delayed in regeneration relative to 40s amputees as well as published timelines for posterior regeneration (Planques et al., 2018). The stages occur ∼1 stage per day, with worms typically reaching stage 5 (late patterning with segment primordia) at 5 DPA (Fig. 5A).

**Figure 5:**
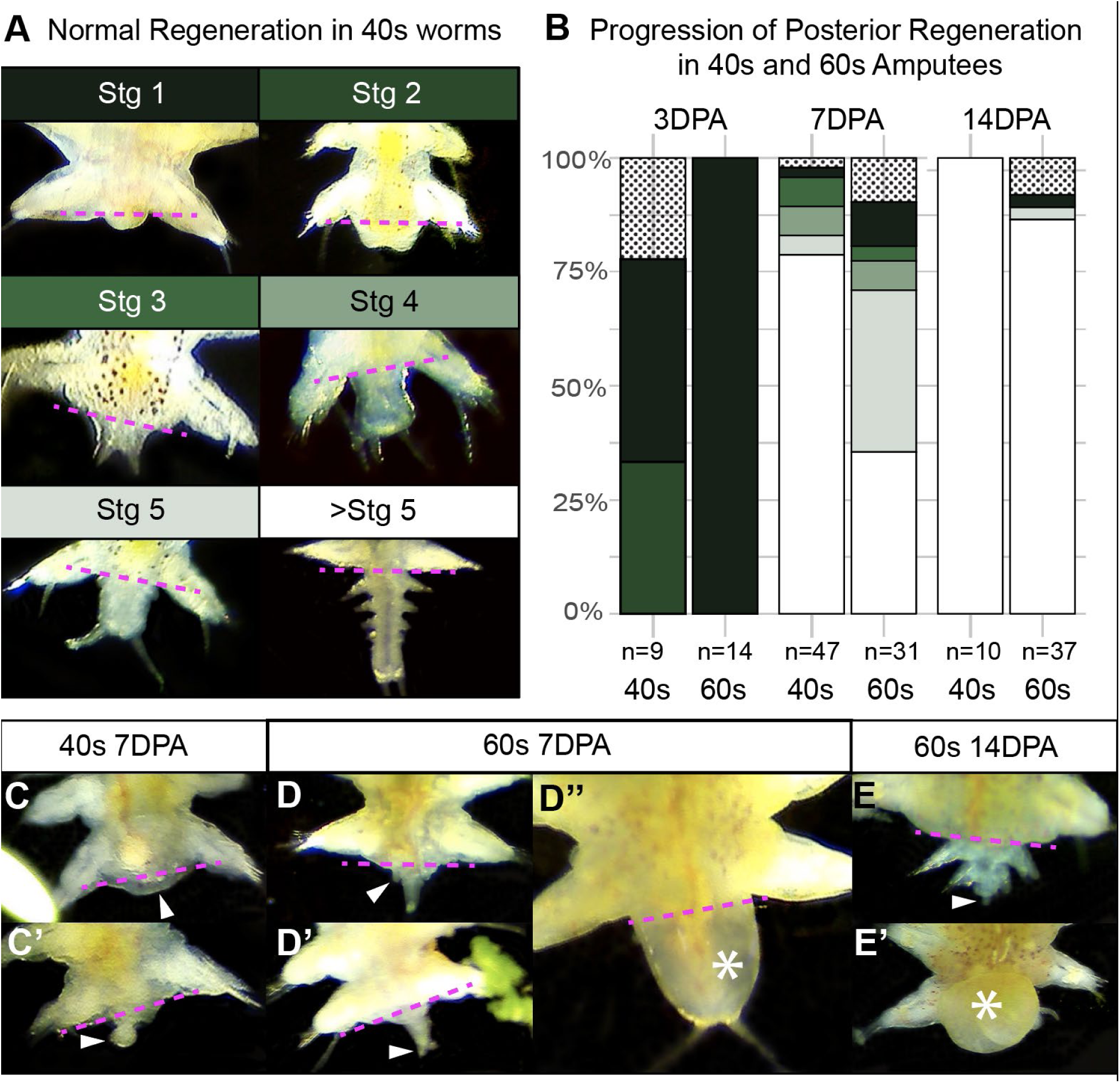
Older Amputees Have Higher Rates of Abnormalities and Delayed Regeneration. A) The stages of normal posterior regeneration in 40s worms. The magenta dashed line indicates the amputation plane separating original tissue (above) and regenerated tissue (below). Stages 1-5 are based on previously published results (Planques et al., 2018): wound healing (Stg 1), formation of the blastema and anus (Stg 2), formation of anal cirri (Stg 3), elongation of the regenerated region and anal cirri (Stg 4), and the patterning of segment primordia (Stg 5). >Stg 5 indicates newly formed segments have visible chaetae, signaling a shift from posterior regeneration into posterior growth. Any morphological deviation was scored as abnormal. B) Bar graphs showing the proportion of each stage at time points 3-14 days post amputation (DPA), color coded by stages in A. Abnormal samples indicated with bars filled with dots. C-E’) Examples of abnormal regeneration in the 40s and 60s worms from 7-14 DPA. The white arrowheads show morphological abnormalities, and white asterisks denote edema. C) Regeneration does not appear to have progressed past wound healing. C’) A small abnormal outgrowth. D) An abnormally small regenerated pygidium that appears to come to a single point rather than symmetrical cirri, with two apparent segment primordia. D’) Anal cirri have formed, but the regenerate is abnormally small. D’’) The regenerated region is swollen and filled with fluid, though patterning appears otherwise correct. E) The pygidium is abnormally small and appears to come to a single point. E’) The area around the wound site is swollen and filled with fluid.

By 7 DPA, over 75% of 40s amputees surpassed stage 5 (chaetae-bearing segments formed) compared to about 40% of 60s amputees . By 14 DPA, 100% of 40s amputees and about 80% of 60s amputees were forming visible segments (Fig. 5B). Abnormalities were more common among 40s amputees than 60s amputees during the early stages of regeneration, at 3 DPA (Fig. 5B). These abnormalities included a failure to form a blastema (Fig. 5C) and aberrant morphology (Fig. 5C’). By 7 DPA, 60s amputees had a slightly higher percentage of abnormalities (Fig. 5B). Aberrant morphology at this stage included improperly scaled and patterned regenerated regions (Fig. 5D-D’) as well as edema (Fig. 5D"). By 14 DPA, no abnormalities remained in 40s amputees (Fig. 5B), one 60s worm had formed chaete-bearing segments but had an improperly formed pygidium (Fig. 5E), and another had edema (Fig. 5E’). All 60s worms that still had abnormal posterior regeneration at 14 DPA died between 19 and 31 DPA.

### Analyses of gonial cluster regeneration in 40s and 60s amputees via HCRs

Our previous experiments established that amputated worms successfully regenerate their gonial clusters and reach sexual maturation without any major differences in terms of fecundity and fertility from unamputated control siblings. Next, we wanted to visualize the early events of gonial cluster regeneration and interrogate age-specific differences using *in situ* Hybridization Chain Reaction (HCR) for the germline marker *vasa*. We focused on the 40s and 60s groups as these two groups had the most pronounced difference from each other in time to maturation. In addition, at the physiological level, these stages highly differ in their state of gametogenesis. 40s worms usually have a variable number of small undifferentiated gonial clusters, while 60s worms have a mix of numerous sexually differentiated clusters nearing maturation as well as some undifferentiated clusters. A minority of 40s worms may have small oocytes due to the asynchronous nature of oogenesis, but there is generally a clear difference in gametogenesis and resource investment between these two ages both in terms of number and maturation stage of the gonial clusters (i.e lower investment in 40s group compared to 60s group). We amputated worms after the 8th segment in both 40s and 60s groups, and fixed samples at 7, 14, or 23 DPA to visualize general *vasa* expression patterns, presence/absence of gonial clusters in the original versus regenerated segments, and the general morphology of the anterior cluster and the gonial clusters.

In both 40s and 60s amputees, gonial clusters appeared in the original segments in the majority of the worms, however the numbers were generally low for the 40s group (median 1, average 2.1) compared to 60s (median 8, average 11.5). At 7 DPA, most 40s amputees had already regenerated new segments, while most 60s amputees were still in mid or late stages of regeneration. Gonial clusters were observed in regenerating and regenerated segments as early as 7 DPA but only 1 gonial cluster in 1 out of 5 in 40s and 2 gonial clusters in 1 out of 6 worms in 60s group (Fig. 6C, D). In both groups, in a small number of samples, we observed gonial clusters right at the border of the original and regenerated segments (Fig. 7B, D, F’). By 14 DPA, the majority of 60s samples (5/6) had gonial clusters (median 3) in the regenerated region, while only one 40s sample (1/5) had 3 gonial clusters in the regenerated region, even though both groups generally had a similar number of segment length (median 20 segments 40s; 17.5 segments 60s). By 23 DPA in both groups the majority of worms had numerous gonial clusters in the regenerated segments (median 3.5 gonial clusters in 40s, 13 in 60s) even though many of the worms had not crossed the 35 segment threshold that is required for normal gonial cluster expansion (median 24.5 segments in 40s, 26.5 segments in 60s). At this stage, 40s amputees still had relatively fewer gonial clusters (median 0.5) in their original segments compared to 60s amputees (median 11) (Fig. 6C, D).

**Figure 6:**
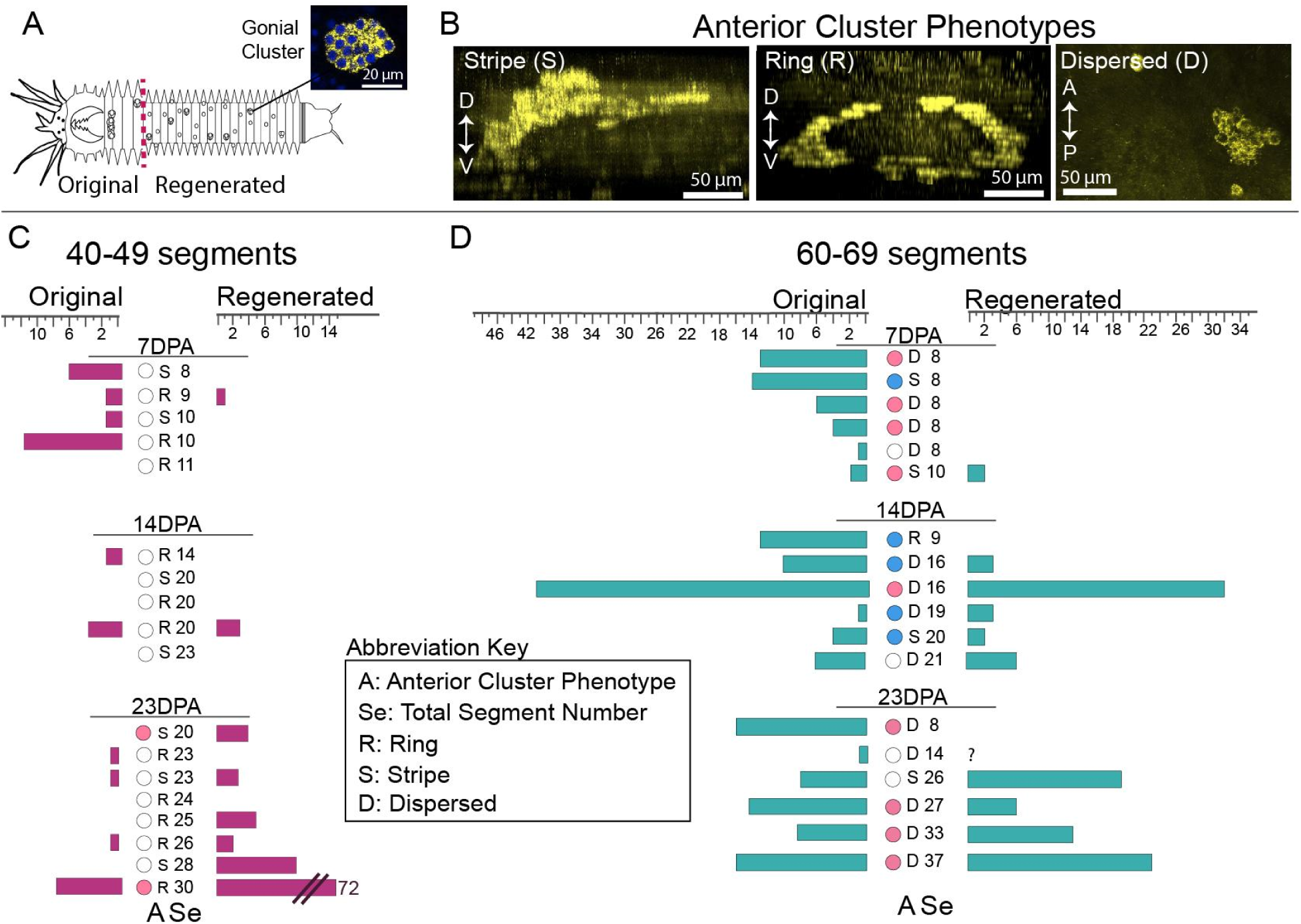
Gonial cluster regeneration across time in the 40s and 60s groups. A) Cartoon showing the regions compared in the graphs. Dashed line indicates the amputation plane. B) Examples of anterior cluster phenotypes visualized via *vasa in situ* HCR (yellow). Note that the first two show dorsal-ventral cross section, where the anterior cluster is either in a stripe layout dorsal to the gut, or it forms a ring that extends dorso-ventrally around the gut. A third phenotype is called dispersed, when a clear pattern is lacking and only smaller clusters of *vasa*-expressing clusters are found in this region. C-D) Graphs showing the number of gonial clusters observed in the 40s or 60s regenerates across time points 7, 14, 23 DPA. Only discreet gonial clusters, and singlet oocytes were included in these counts (see Fig. 7G for examples) . The circles indicate whether observed clusters have started sexual differentiation (blue/pink for M/F, empty for undifferentiated or NA, if there are no clusters observed). The “A” column indicates the state of the anterior cluster: “D” for dispersed, “R” for ring, “S” for stripe. The column “Se” reports the total number of segments in each sample. Samples are arranged from shortest to longest in terms of segment number.

**Figure 7:**
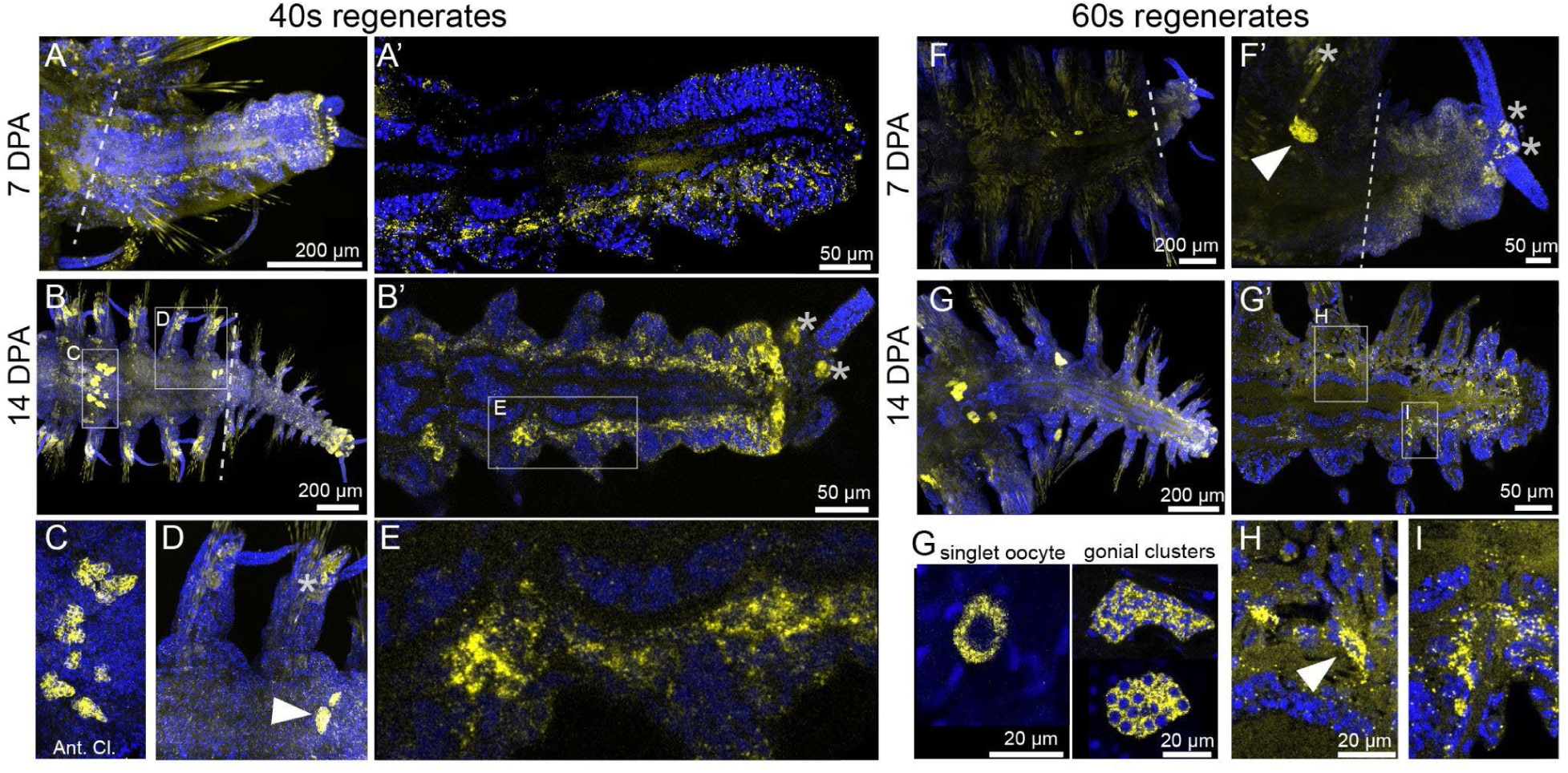
*in situ* HCR for *vasa* expression in 40s and 60s regenerates. Examples of samples from Figure 6 shown. At 7 DPA (A-A’) and 14 DPA (B-B’) in 40s worms, there is a broad *vasa* expression (yellow) that looks distinct from gonial clusters (arrowheads in D and F’). E) Zoomed version of the broad expression in the 40s amputee. This broad expression is also present in 60s regenerates but more prominent at 14 DPA (G-G’). H and I show zoomed versions of the *vasa*-expressing cells that are found during regeneration. Compared with the singlet oocytes or gonial clusters, these cells appear less organized, have smaller nuclei, and are generally located in the region between the gut and the epidermis. Zoomed version of the anterior cluster (example of “stripe”) is shown in C. Blue color is DAPI nuclear stain. Dashed line indicates the amputation plane. Stars indicate non-specific signal that is commonly found in chaetal sacs, chaetae and the pygidium (B’, D, F’).

Next, we wanted to assess changes in the anterior cluster (AC) during regeneration in either group. The AC starts forming when the primordial germ cells migrate anteriorly in larval stages, and as the worms grow longer the AC forms (presumably from the proliferation of the germ cells) between the 5th and 6th segments, around the pharynx (Kuehn et al., 2021; Rebscher et al., 2012, 2007). The morphology of the AC changes as the worms add segments, and these morphological phenotypes are classified as follows (Fig. 6B): ‘ring’, where the clusters are arranged in a ring shape around the gut/pharynx with a clear ventral and dorsal component; ‘stripe’, where the clusters are arranged linearly, usually dorsally; and ‘dispersed’, where there is not a notable organization of an anterior cluster in this specific region, although there may be some individual clusters nearby in no particular pattern. When scored, 40s and 60s amputees showed strikingly different AC phenotypes (Fig. 6C, D). For 40s, all samples at all timepoints had a distinct anterior cluster (18/18; Fig. 6C, 7C). Many samples had a clear ring phenotype (11/18), with a few samples with the stripe phenotype (7/18). There did not appear to be a trajectory towards ring or stripe phenotypes as regeneration progressed, and there were no samples with the dispersed AC phenotype. In contrast, 60s AC phenotypes were dramatically different: in most 60s amputees, the anterior cluster was dispersed (13/18; Fig. 6D). This observation is consistent with anecdotal experience in the lab in uninjured worms over 60 segments, we often observe no AC (Kuehn et al., 2021).

In addition to gonial clusters, we noticed that in both 40s and 60s groups, there was a broad *vasa* expression in the newly-regenerated segments in the region between the gut and the epidermis (Fig. 7A’, B’, E, G’). This pattern was most pronounced at 14 DPA in both groups, but it arose as early as 7 DPA in the 40s amputees (60s amputees were still in the regeneration phases at this time point). Furthermore, in both 40s and 60s samples we began to see single cells or small groups of cells expressing *vasa*, but their overall morphology was distinct from gonial clusters or singlet oocytes (Fig. 7G, H, I). These putative progenitor cells were first observed at 14 DPA and persisted to 23 DPA when the broad *vasa* expression was no longer present except for in the posterior growth zone. These putative progenitor cells were typically in the areas surrounding the gut, the body wall musculature, or parapodia, but not in the coelom. They appeared smaller than gonial cluster cells, with smaller and less round nuclei.

## DISCUSSION

### The effect of regeneration on reproduction, growth and survival

Many annelids, including *P. dumerilii* can regenerate germ cells and gonads (Özpolat, 2023). While there have been anecdotal notes on *P. dumerilii’*s ability to regenerate its germline during posterior regeneration (Hauenschild and Fischer, 1969; Rebscher, 2014), systematic studies on this process to see whether removal and regeneration of germ cells have an impact on fertility and fecundity have not been carried out before. In some annelids injury can negatively impact reproduction, measured by the production of gametes, or gamete, cocoon, and brood numbers (Gibbs, 1968; Hill et al., 1982; Rennolds and Bely, 2023; Zajac, 1985). To address this in *P. dumerilii*, we implemented the most severe survivable amputation at three different developmental stages with differential germline investment states. The oldest worms (60s) had extensive germline investment (late stages of gametogenesis, e.g. with vitellogenic oocytes) and were on the cusp of sexual metamorphosis, while the younger worms (40s or 50s) had only undifferentiated germ cells, or were at the early stages of gametogenesis, respectively. We expected that older worms that have high gametogenesis investment potentially could show higher rates of adverse effects on fecundity upon injury and regeneration compared with younger worms. However, we found that all 40s, 50s, and 60s amputees which regenerated have normal fertility and fecundity, measured by the number of eggs released, percentage of eggs fertilized, and successful maturation of the next generation.

While reproductive success was not overall different in the 3 groups, we observed differences in time to maturation. Here, we expected that the older group would be more delayed in the time to maturation due to the larger loss experienced, however we found the opposite to be the case: 60s amputees reached maturation earlier than 50s and 40s amputees. A simple explanation for this could be the overall growth rate differences. As previously shown, number of segments is a good metric of growth in annelids, and especially in *P. dumerilii*, worms are known to have to reach a particular segment number threshold for developmental transitions (Kuehn et al., 2021). Therefore we reasoned that even though all amputees are reduced to a total of 8 segments, older worms may be growing faster than younger worms, hitting the developmental thresholds faster, and therefore reaching sexual maturation faster. Indeed, we found that older 60s amputees had an overall higher segment addition rate compared with younger 40s amputees. However, the growth rate difference was not drastic enough to be the only factor explaining the difference in time to maturation. Other factors in the anterior 8 segments remaining after amputation could be physiological, morphological, and molecular changes that occur in *P. dumerilii* before the visible onset of maturation (Dahlitz et al., 2023; Fischer et al., 2010; Fischer and Hoeger, 1993; Schenk et al., 2019, 2016). For example, there may be differential storage of nutrients, including lipids, that could enable the rapid growth observed in older amputated worms. The older worms tend to also get larger and they may be more efficient in feeding. The neurohormonal state is also likely to play a major role, as with age, complex hormonal shifts take place (see below for details). Overall, due to these changes, older worms may be primed to return to the metabolic, hormonal states associated with maturation compared to younger worms, therefore reaching maturation faster.

We also observed, however, that amputation and regeneration did not come without a cost for the 60s amputees, which had a higher rate of mortality or abnormal regeneration after amputation, relative to the 40s and 50s amputees. That is, if 60s amputees could successfully regenerate the posterior body axis they also regenerated germ cells without an adverse effect on their reproductive success, but on the other hand some samples failed regeneration altogether, or had higher mortality even after having regenerated segments but had stalled growth. One particular 60s amputee never grew beyond 16 segments, eventually died and upon investigation had several large oocytes in the regenerated segments. As gametes mature they are known to signal negative feedback to suppress growth and regeneration, so the worms can initiate sexual maturation. The failure of regeneration in some of the 60s amputees could be due to the hormonal state differences and these signals from maturing gametes present in the original segments (see below for detailed discussion on the hormones).

Finally, injury can lead to delays in developmental transitions in some organisms. In *Drosophila*, injury to the imaginal disc during the regenerative competent larval phase results in a developmental delay to accommodate regeneration (Jaszczak and Halme, 2016; Karanja et al., 2022). In annelids, several works suggested that injury affects reproduction (Gibbs, 1968; Hill et al., 1982; Rennolds and Bely, 2023; Zajac, 1985). In *Scoloplos*, Gibbs collected wild worms and observed that many of the worms were injured and regenerating, or had regenerated at some point. Some worms that appeared to have several regeneration cycles were delayed in their production of gametes, and/or produced fewer cocoons as a result. Therefore injury negatively affected fecundity/reproduction in this species (which is not semelparous). In *Polydora*, Zajac made amputations to cause injury (but did not remove segments that contained the gametes) and showed that injury negatively affected fecundity (measured in terms of number of eggs or brood size). In addition, posteriorly amputated worms took approximately twice as long to the production of the first brood and produced less broods compared to the controls. In our experiments, when we compared control unamputated siblings with the amputated groups, we found that amputation and regeneration did not cause a delay to reach sexual maturation in younger groups (40s, 50s) although the older group (60s) was delayed to maturation compared to their controls. This lack of delay in the younger groups was surprising because amputees need to recover dozens of lost segments and regenerate germ cells, but this setback does not come with any significant delay to maturation. One possibility that may explain this observation is faster growth (segment addition) after regeneration. This phenomenon has been observed in *P. dumerilii* before where segment addition rate is highly increased in worms that have recently regenerated (Gazave et al., 2013). Indeed our growth curves also support this: we observe that segment addition rates spiked following amputation, remained high until ∼30 DPA (regeneration is completed between 5-10 DPA), and eventually appeared to plateau and approach control levels by 63 DPA. Therefore, we see an overall increased growth rate in all amputated groups compared with the control siblings.

### Hormonal control of growth and regeneration

It is possible complex hormonal interactions and changes explain some of the observations outlined above. The changes in the brain’s hormonal composition has been studied extensively in *P. dumerilii* and other Nereididae, and the hormonal states are known to change with growth, age, and during injury (Andreatta et al., 2020; Clark, 1969; Clark and Scully, 1964; Golding, 1974, 1967a, 1967b; Hauenschild, 1974, 1966; Hofmann, 1976; Lawrence and Soame, 2009; Olive and Clark, 1978; Scully, 1964). The sesquiterpenoid methylfarnesoate (MF) was recently identified as the ’brain hormone’ in *P. dumerilii* (Schenk et al., 2016). MF is known to promote growth and regeneration, inhibit sexual maturation, and mediate developmental transitions in *P. dumerilii* and other annelids (Álvarez-Campos et al., 2023; Biggers and Laufer, 1999, 1996; Hauenschild, 1974, 1966; Hofmann, 1976; Lawrence and Soame, 2009). Brain transplantation experiments have demonstrated that the brain hormone activity generally decreases with age (Hofmann, 1976; Scully, 1964), though the exact trajectory of hormone levels over the worm’s lifetime remains unclear. The decreasing levels of MF with age could explain the higher mortality and abnormal regeneration observed in 60s amputees. Other transplantation experiments suggested that brain hormone activity increases in response to amputation (Golding, 1967a), which is thought to be one explanation for faster growth and segment addition rates in recently-regenerated worms. However, no direct measurements of MF have been carried out in these contexts yet, so it remains to be shown whether these observations directly correlate with changes in MF hormone levels.

Another possible hormonal factor is thought to exert a negative feedback on the MF activity, triggering sexual metamorphosis and causing the loss of regenerative competency in *P. dumerilii* and other Nereididae (Andries, 2001; Porchet and Cardon, 1976; Schenk et al., 2016). The identity of this compound is still unknown but several lines of evidence suggest that this ‘feedback substance’ is produced by the maturing gametes, eventually inhibiting the MF activity altogether, leading to sexual maturation. Transplantation of oocytes, lysates from the maturing gametes, or injection of coelomic contents were shown to exert negative feedback on the activity of the brain hormone (Andries, 2001; Durchon, 1952; Hofmann, 1975; Porchet, 1967; Porchet and Cardon, 1976; Porchet and Durchon, 1968). In *Nereis diversicolor*, worms with oocytes larger than 140 µm show diminished posterior regeneration and worms with oocytes exceeding 180 µm can not regenerate at all (Golding, 1967b).

The opposing effects of the brain hormone and the feedback substance could explain some of our results, which may illustrate the trade-offs between germline and somatic regeneration under this hormonal regulatory system. In our experiments, while 60s amputees were capable of full germline regeneration and matured earlier than younger amputees, they also showed higher rates of abnormal regeneration and mortality, with one early wave of deaths by 7 DPA (the completion of posterior regeneration) and several more deaths scattered over the rest of the experiment. Our amputations after segment 8 remove most of the gonial clusters but do not remove the anterior cluster or any gonial clusters remaining in the 8 head segments. The 60s group typically has gametes closer to maturation, and a dispersed anterior cluster which is effectively turning into gonial clusters going into gametogenesis. In contrast, in most 40s samples, gonial clusters were not progressed enough to identify whether they were becoming oocytes or sperm. Therefore the 60s group could presumably have gonial clusters remaining in the 8 original segments with a high negative feedback effect on the brain hormone activity. It is possible that large oocytes, in particular, could interfere with regeneration, possibly through MF inhibition. This could contribute particularly to the mortality that occurs in worms that have successfully added segments before arresting growth and then dying, as observed in one sample that we fixed on the threshold of death, and found oocytes that appeared to have mature size and morphology. It is important to note that several other factors, including mechanical failures of wound closure, could also contribute to increased mortality. Future studies will be required to quantify MF levels following amputation at different time points in different segment size groups to assess the relationship to segment addition rates and mortality.

### Developmental thresholds

Organisms have mechanisms to assess growth and nutritional state to make resource allocation decisions towards more growth or sexual metamorphosis and reproduction (Henry and Hart, 2005; Hyun, 2018; Lui and Baron, 2011; Olive and Clark, 1978). For example, holometabolous insects need to reach a threshold body weight for metamorphosis (Tennessen and Thummel, 2011). Studies of polychaete reproductive physiology and endocrinology indicate that in some species maturity, oogenesis, and/or spawning are arrested or delayed until regeneration is complete following tissue loss, usually anterior segments which produce hormones controlling these facets of polychaete reproduction (Olive and Clark, 1978). Similarly in *P. dumerilii*, our previous studies demonstrated a threshold of about 35 segments when small germline clusters start populating the body cavity (germline expansion) and that if worms are amputated before germline expansion, they must still reach ∼35 segments to undergo germline expansion. However, in the current study, we observe that once worms have reached and passed the germline expansion stage, they do not need to grow back to 35 segments to start producing gonial clusters after regeneration. For example, by 23 DPA, worms have numerous gonial clusters even though most of them have not reached the ∼35 segment threshold: 40s amputees that have started regenerating gonial clusters range between 20-30 segments, and 60s amputees that have started regenerating gonial clusters range between 26-37 segments. These results suggest that once worms have crossed the ∼35-segment milestone, they no longer need to reach 35 segments to produce gonial clusters again.

### Potential sources of regenerated gonial clusters and future directions

Many annelids express germline/multipotency genes such as *vasa*, *piwi*, *nanos* in tissues that are not strictly germline (Gazave et al., 2013; Kostyuchenko and Smirnova, 2023; Özpolat and Bely, 2016, 2015; Planques et al., 2018). For example, in *P. dumerilii*, *vasa* is expressed not only in the gonial tissues, but also in the posterior growth zone, and in the regeneration blastema. It is still unclear if these tissues are able to give rise to both somatic and germ cells. In this study, we also found *vasa* expression to persist days and weeks after regeneration is completed in a broad pattern that extends beyond the growth zone into many segments. This expression appears very different from expression patterns observed in worms that were amputated in stages before gonial clusters form (e.g. 10s or 20s worms) (Gazave et al., 2013; Kuehn et al., 2021; Planques et al., 2018). These worms have individual or groups of *vasa*-expressing cells that do not look like either the singlet oocytes, typical gonial clusters, or the posterior growth zone expression that is typically confined to the segment addition zone and a few newly-produced segments. Therefore these cells could be a potential source of regenerated cell types, including the gonial clusters. Future studies with genetic lineage tracing will be able to directly address this open question.

In addition, due to the unique biology and morphology of *P. dumerilii*, where germline clusters float freely throughout the coelom, we could not determine whether the clusters that we observed in the newly regenerated segments, especially at the earlier regeneration time points, were themselves regenerated *in situ*, or were a product of the original gonial clusters that drifted into the regenerated region from the original segments, or both. It is important to note that especially in 60s group, gonial clusters are generally progressed in their gametogenesis, and if new gonial clusters are made from these existing gonial clusters, how do progressed gonial clusters at such distinct states of gametogenesis give rise to new gonial clusters will be an interesting area of inquiry. It is possible some gonial clusters regress back to an earlier more undifferentiated stage to undergo mitosis. In addition, if the new gonial clusters only regenerate from the original anterior cluster and/or gonial clusters, it will be interesting to investigate whether these sources can be depleted upon repeated amputation-regeneration cycles. In previous studies we observed that the anterior cluster eventually disperses in control worms, and in this study, we also show that it disperses in 60s amputees. The worms with the dispersed anterior cluster phenotype may respond to the repeated amputation-regeneration challenge differently. Future studies focusing on genetic lineage tracing, or live cell tracing using tissue-specific expression of fluorescent markers will address these open questions.

## ACKNOWLEDGEMENTS

We thank the Özpolat Lab members for their helpful feedback.

---

**Figure S1:**
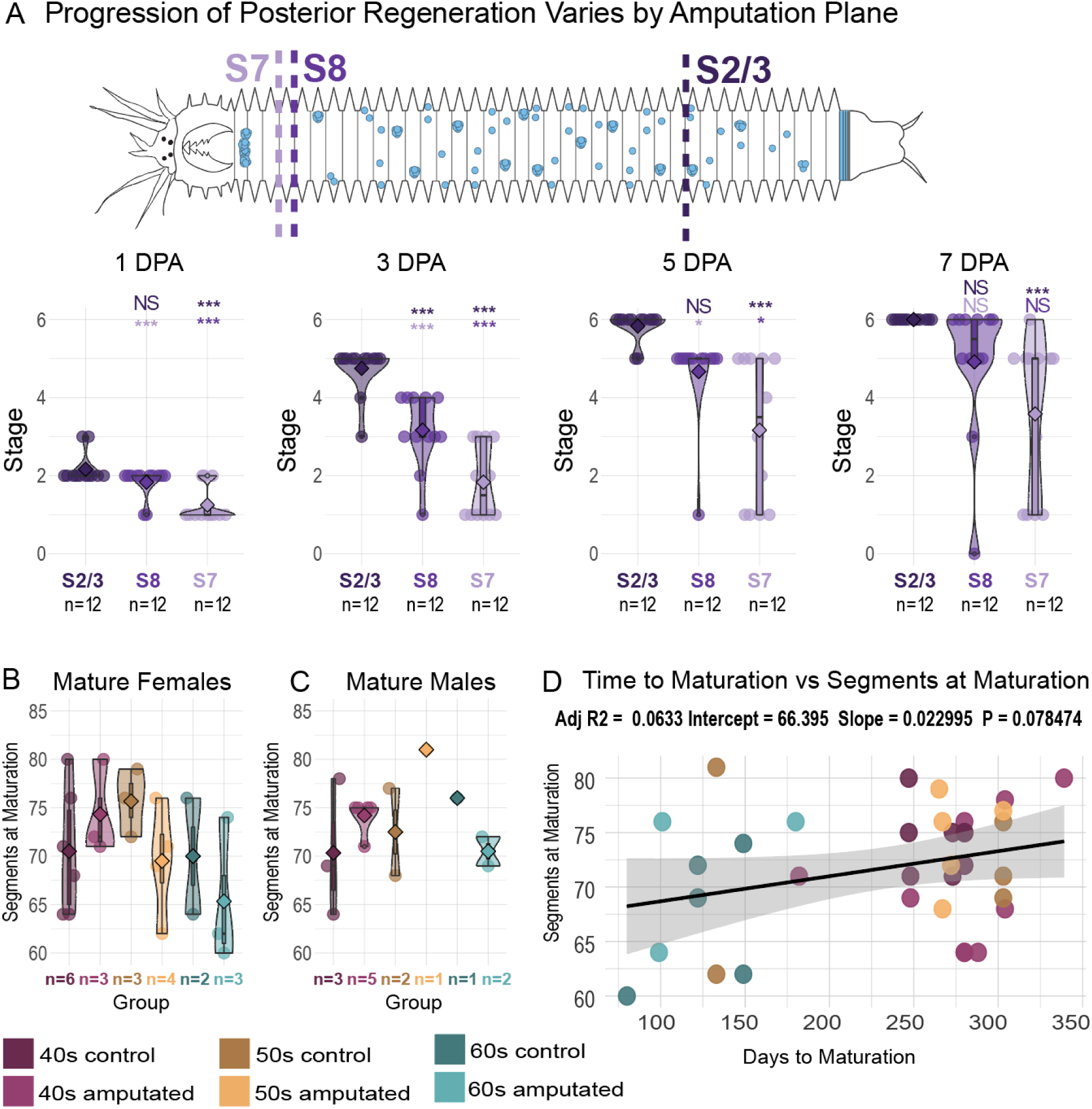
Worms cut at different locations show variable progression of posterior regeneration. Amputations after the 7th pair of chaetae, the 8th pair of chaetae, or approximately 2/3rds down the length of the body (removing ∼1/3rd) were compared. The stage of regeneration was recorded for each group at A) 1 day post amputation (DPA), B) 3 DPA, C) 5 DPA, and D) 7 DPA. By 7 DPA, worms amputated after the 8th pair of chaetae did not significantly differ from worms with ∼1/3rd of segments removed (t test, p=0.058). Therefore the 8th segment was chosen as the amputation plane to maximize segment removal for subsequent experiments. E,F) The number of segments at maturation showed no significant differences between groups when separated by sex. G) When all groups are plotted by days to maturation against the number of segments at maturation, there is not a statistically significant linear correlation (p=0.08).

**Figure S2.**
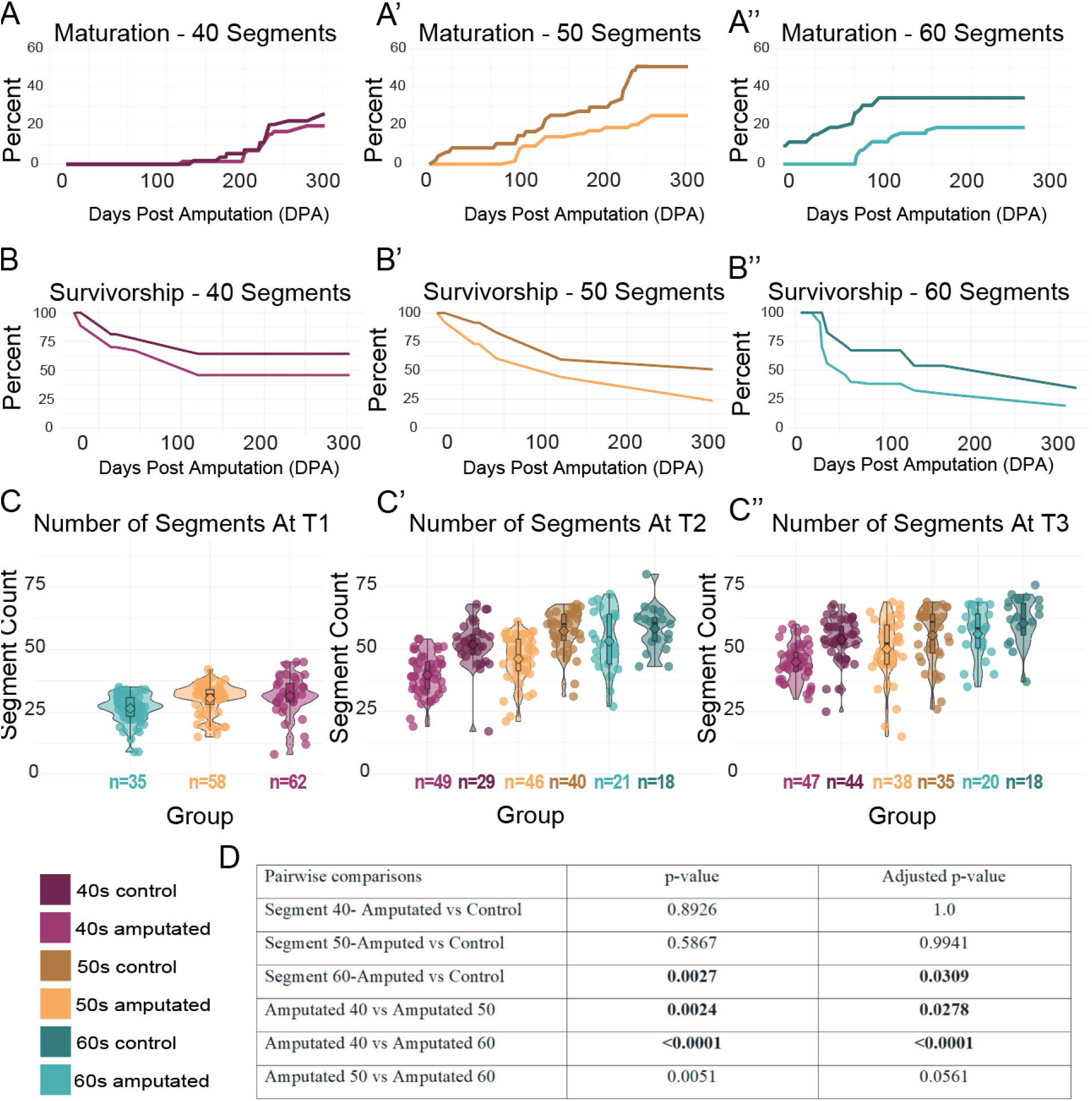
A-A’’) Plots show maturation over time for three segment groups. B-B’’) Plots show survivorship over time for three segment groups. Color codes for all groups are indicated in the lower left. C-C’’) The number of segments at T1 (28 days post amputation (DPA)). Only amputated groups represented in this count. C’) The number of segments at T2, 63 DPA. C’’) The number of segments at T3, 84 DPA. Both amputated and control groups are shown in C’ and C’’. D) P values for pairwise comparisons of time to maturation before and after Tukey-Kramer adjustment. Statistically significant values in bold (p>=0.05).

**Figure S3.**
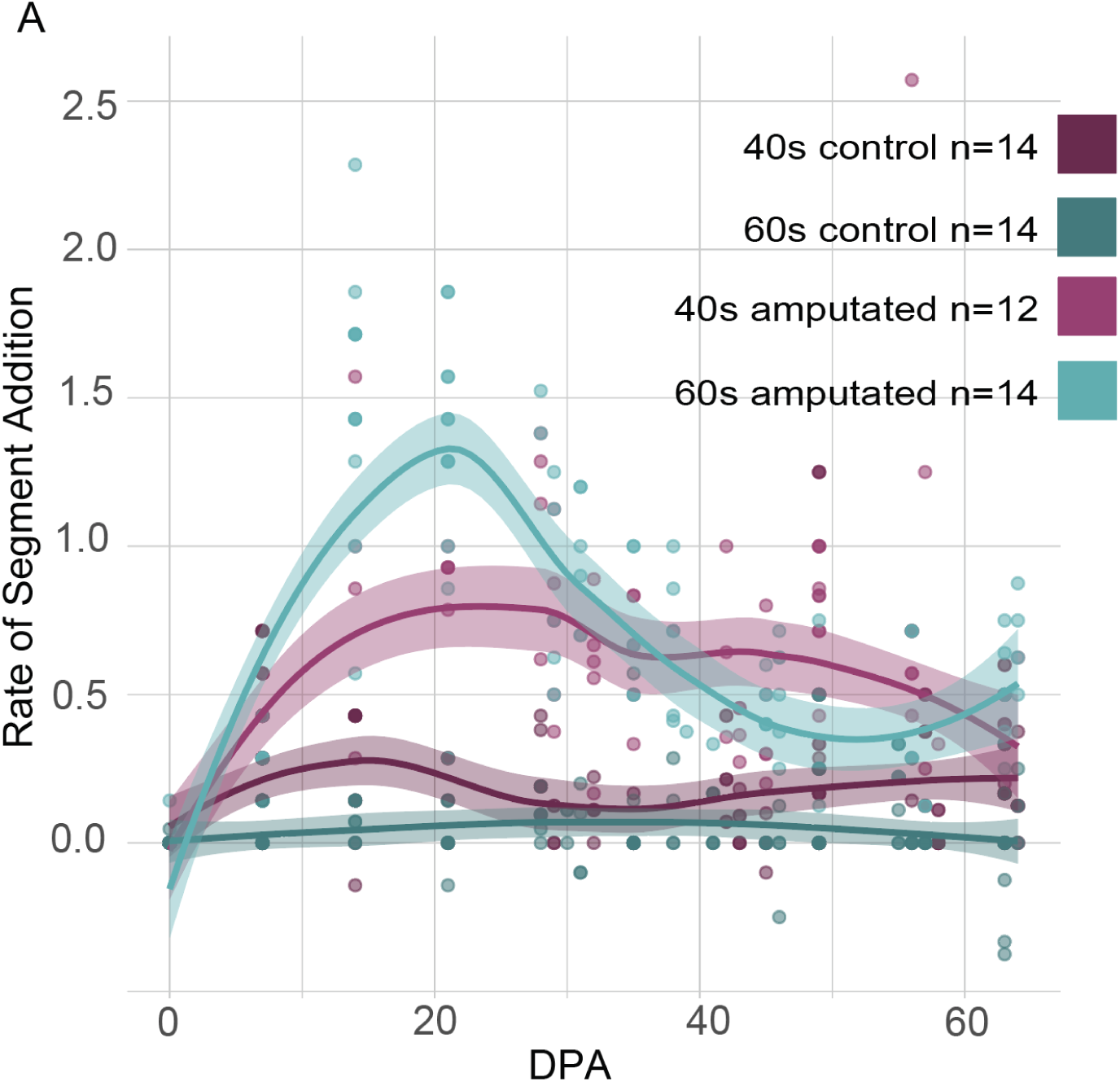
A) Worms were counted approximately once per week. Segment addition rate was calculated as the number of segments added divided by the number of days since the last measurement. Only worms that survived until the final measurement time point at 63 DPA were included in this analysis, since worms that die slow down and then stop adding segments in advance of their death and the elevated mortality rate of 60s amputees would obscure trends in segment addition rate. A few individual worms have negative growth rates; this is likely due to +/-1 variation in segment counting and occurs exclusively in controls with slow to no segment addition, so +/-1 results in a negative rate.

