## Supplementary File 01 for "The cost and payout of age on germline regeneration and sexual maturation in *Platynereis dumerilii*"

### Supplementary Files for R codes

#### amp_pilot.csv (Suppl. Fig. 1)

This spreadsheet contains the stage of posterior regeneration over time in worms with three different amputation sites.

#### AE.csv (Fig. 1, Suppl. Fig. 1)

This spreadsheet contains several endpoints for maturation: length at maturation, the number of days it took to reach maturation, and the location of the atokous/epitokous border. Other variables include the initial segment group, whether or not the worm was amputated, the sex of the mature worm, and which box the worm was cultured in.

#### form_eggs.csv (Fig. 1)

This spreadsheet contains fecundity data for each mating assessed, including the number of eggs released and the percentage fertilized.

#### format.csv (Fig. 2)

This spreadsheet contains the cumulative number of mature animals over time per condition in the experiment tracking time to maturation. Other variables include environmental/culturing variables regarding the room temperature and the maturation cycle.

#### format_noncumulative.csv (Fig. 2, Suppl. Fig. 2)

This spreadsheet contains the number of mature worms found on a given day for the experiment tracking time to maturation. Same data as format.csv, but noncumulative.

#### death_cumulative.csv (Suppl. Fig. 2)

This spreadsheet contains the cumulative death totals over time for the experiment tracking time to maturation. Other variables include the initial segment group and whether or not the worm was amputated. Shows per treatment/condition, boxes pooled.

#### AF_4wk_scounts.csv (Suppl. Fig. 2)

This spreadsheet contains the segment count data at 28 DPA for the experiments primarily concerned with the time to maturation. These are not associated with individual worms.

#### AF_8wk_scounts.csv (Suppl. Fig. 2)

This spreadsheet contains the segment count data at 63 DPA for the experiments primarily concerned with the time to maturation. These are not associated with individual worms.

#### AF_12wk_scounts.csv (Suppl. Fig. 2)

This spreadsheet contains the segment count data at 84 DPA for the experiments primarily concerned with the time to maturation. These are not associated with individual worms.

#### singles.csv (Fig. 3, Fig. 4, Fig. 5)

This spreadsheet contains the segment counts over time for individual worms. Other variables include days surviving (if died), original segment number prior to amputation, and stage of blastema during posterior regeneration.

#### t4_rates.csv (Suppl. Fig. 3)

This spreadsheet contains the rates of growth associated with individual worms from singles.csv. The rate assigned to a timepoint was calculated by the number of segments added divided by the number of days since the previous segment count.
