## Supplementary File 02 for "The cost and payout of age on germline regeneration and sexual maturation in *Platynereis dumerilii*": Metzger_Fig1.R

```
#Figure 1:

#Loading packages (used across all figures)
library(ggplot2)
library(gcookbook)
library(dplyr)
library(hrbrthemes)
library(ggpubr)
library(ggsignif)
library(tidyr)
library(tidyverse)
library(reshape)
library(ggbeeswarm)
hrbrthemes::import_roboto_condensed()

#Setting color schemes
colors = c("#71254F","#a23671", "#B2793D", "#ffae58", "#317B7D", "#47B1B4", "#2F6A3A", "#93CF5B")

#Set working directory
setwd("~/Documents/Ozpolat/Experiments/Amputation & Fecundity")

#Reading in data
form_eggs = read.csv(file="form_eggs.csv", header=T, sep=",")
AE = read.csv(file="AE2.csv", header=T, sep=",")

#Fig 1E
ggplot(AE, aes(x=as.factor(Index), y=Segments)) + 
  ylim(55,85) + xlab("Initial Segment Group") + 
  ylab("Segment Count At maturation") + geom_violin(adjust=0.5, alpha=0.5, aes(fill=Index)) + 
  geom_boxplot(width=0.1, outlier.size=0, outlier.shape=21, aes(fill=Index)) + 
  stat_summary(fun="mean", geom="point", shape=23, size=5, aes(fill=Index))+ 
  geom_quasirandom(size=5, alpha=0.7, aes(color=Index)) + 
  theme_ipsum(axis_title_just = 0, axis_title_size =12) + 
  scale_fill_manual(values=colors) + 
  scale_color_manual(values=colors) + 
  guides(fill=FALSE, color=FALSE)
shapiro.test(AE$Segments) #yes, W = 0.95119, p-value = 0.1233
length_mat_aov = aov(Segments ~ Group, data = AE)
summary(length_mat_aov)

length(AE$Date[which(AE$Group=="s40_control")]) #10
length(AE$Date[which(AE$Group=="s40_amputated")]) #8
length(AE$Date[which(AE$Group=="s50_control")]) #5
length(AE$Date[which(AE$Group=="s50_amputated")]) #6
length(AE$Date[which(AE$Group=="s60_control")]) #3
length(AE$Date[which(AE$Group=="s60_amputated")]) #6

#Fig 1E alternative: Pooled
ggplot(AE, aes(x=as.factor(Group1), y=Segments)) + 
  ylim(55,85) + xlab("Initial Segment Group") + 
  ylab("Segment Count At maturation") + 
  geom_violin(adjust=0.5, alpha=0.5, aes(fill=Group1)) + 
  geom_boxplot(width=0.1, outlier.size=0, outlier.shape=21, aes(fill=Group1)) + 
  stat_summary(fun="mean", geom="point", shape=23, size=5, aes(fill=Group1)) + 
  geom_quasirandom(size=5, alpha=0.7, aes(color=Group1)) + 
  theme_ipsum(axis_title_just = 0, axis_title_size =12) + 
  scale_fill_manual(values=colors[c(8,7)]) + 
  scale_color_manual(values=colors[c(8,7)]) + 
  guides(fill=FALSE, color=FALSE)
length(AE$Date[which(AE$Group1=="amputated")]) #20
length(AE$Date[which(AE$Group1=="control")]) #18

#Atokous/epitokous border
#n's
length(AE$Date[which(AE$Group1=="amputated" & AE$Sex=="Female")]) #12
length(AE$Date[which(AE$Group1=="amputated" & AE$Sex=="Male")]) #8
length(AE$Date[which(AE$Group1=="control" & AE$Sex=="Female")]) #12
length(AE$Date[which(AE$Group1=="control" & AE$Sex=="Male")]) #6

#Fig 1G: females
AE_female = subset(AE, (Sex %in% c("Female")))
ggplot(AE_female, aes(x=as.factor(Group1), y=A.E.border)) + 
  xlab("Amputated vs Control") + ylab("A/E Segment Boundary") + 
  geom_violin(adjust=0.5, alpha=0.5, aes(fill=Group1)) + 
  geom_boxplot(alpha=1, aes(fill=Group1), width=0.1, outlier.size=0, outlier.shape=21) + 
  stat_summary(alpha=1, aes(fill=Group1), fun="mean", geom="point", shape=23, size=5) + 
  geom_quasirandom(size=5, alpha=0.7, aes(color=Group1)) + 
  theme_ipsum(axis_title_just = 0, axis_title_size =12) + 
  scale_color_manual(values=colors[c(8,7)]) + 
  scale_fill_manual(values=colors[c(8,7)]) + 
  guides(fill=FALSE, color=FALSE)

#Fig 1H: males
AE_male = subset(AE, (Sex %in% c("Male")))
ggplot(AE_male, aes(x=as.factor(Group1), y=A.E.border)) + 
  xlab("Amputated vs Control") + 
  ylab("A/E Segment Boundary") + 
  geom_violin(adjust=0.5, alpha=0.5, aes(fill=Group1)) + 
  geom_boxplot(alpha=1, aes(fill=Group1), width=0.1, outlier.size=0, outlier.shape=21) + 
  stat_summary(alpha=1, aes(fill=Group1), fun="mean", geom="point", shape=23, size=5) + 
  geom_quasirandom(size=5, alpha=0.7, aes(color=Group1)) +
  theme_ipsum(axis_title_just = 0, axis_title_size =12) + 
  scale_color_manual(values=colors[c(8,7)]) + 
  scale_fill_manual(values=colors[c(8,7)]) + 
  guides(fill=FALSE, color=FALSE)

#Fig 1I:
ggplot(form_eggs[which(form_eggs$Mature.sex=="Female"),], aes(x=as.factor(T), y=Total)) + 
  xlab("Amputated vs Control") + ylab("Number of eggs per fertilization") + 
  geom_violin(adjust=0.5, alpha=0.5, aes(fill=T)) + 
  geom_boxplot(alpha=1, aes(fill=T), width=0.1, outlier.size=0, outlier.shape=21) + 
  stat_summary(alpha=1, aes(fill=T), fun="mean", geom="point", shape=23, size=5) + 
  geom_quasirandom(size=5, alpha=0.7, aes(color=T)) +
  theme_ipsum(axis_title_just = 0, axis_title_size =12) + 
  scale_color_manual(values=colors[c(8,7)]) + 
  scale_fill_manual(values=colors[c(8,7)]) + 
  guides(fill=FALSE, color=FALSE)
#n's
length(form_eggs$Total[which(form_eggs$T=="amputated" & form_eggs$Mature.sex=="Female")]) # 6
length(form_eggs$Total[which(form_eggs$T=="control" & form_eggs$Mature.sex=="Female")]) #9

#significance test
form_eggs_f = form_eggs[which(form_eggs$Mature.sex=="Female"),]
wilcox.test(form_eggs_f$Total[which(form_eggs_f$T=="amputated")],form_eggs_f$Total[which(form_eggs_f$T=="control")]) #p-value = 0.8596 NS

#Fig 1J:
ggplot(form_eggs, aes(x=as.factor(T), y=X..Fert)) + 
  xlab("Amputated vs Control") + 
  ylab("Percent Fertilized") + 
  geom_violin(adjust=0.5, alpha=0.5, aes(fill=T)) + 
  geom_boxplot(alpha=1, aes(fill=T), width=0.1, outlier.size=0, outlier.shape=21) + 
  stat_summary(alpha=1, aes(fill=T), fun="mean", geom="point", shape=23, size=5) + 
  geom_quasirandom(size=5, alpha=0.7, aes(color=T)) +
  theme_ipsum(axis_title_just = 0, axis_title_size =12) + 
  scale_color_manual(values=colors[c(8,7)]) + scale_fill_manual(values=colors[c(8,7)]) + 
  guides(fill=FALSE, color=FALSE)

#n's
length(form_eggs$Total[which(form_eggs$T=="amputated")]) # 7
length(form_eggs$Total[which(form_eggs$T=="control")]) #13

#significance test
wilcox.test(form_eggs_f$X..Fert[which(form_eggs_f$T=="amputated")],form_eggs_f$X..Fert[which(form_eggs_f$T=="control")]) #p-value = 0.4787 NS
```
