## Supplementary File 02 for "The cost and payout of age on germline regeneration and sexual maturation in *Platynereis dumerilii*": Metzger_Fig2.R

```
#Figure 2:

#Loading packages (used across all figures)
library(ggplot2)
library(gcookbook)
library(dplyr)
library(hrbrthemes)
library(ggpubr)
library(ggsignif)
library(tidyr)
library(tidyverse)
library(reshape)
library(ggbeeswarm)
hrbrthemes::import_roboto_condensed()

#Setting color schemes
colors = c("#71254F","#a23671", "#B2793D", "#ffae58", "#317B7D", "#47B1B4", "#2F6A3A", "#93CF5B")

#Set working directory
setwd("~/Documents/Ozpolat/Experiments/Amputation & Fecundity")

#Reading in data
format = read.csv("forMAT-final.csv", header=T) #cumulative maturations, not separated by box
format_nc = read.csv(file="format_noncumulative.csv", header=T, sep=",")
#untabling it
expanded_T = untable(format_nc[,c(1:8)], num=format_nc[,9])[,c(1:6)] #does not preserve information of sex, but sex was not significant by univariate ANOVA so alright for graphing

#Fig2B: Observed
format_amp = subset(format, Condition == "AMP")
ggplot(format_amp, aes(x = DPA, y = Percent_M, fill = as.factor(Index), colour=Index)) + 
  ylim(0, 60) + xlab("Days Post Amputation (DPA)") + ylab("Percent Mature") + 
  geom_line(size=1.5) + 
  theme_ipsum(axis_title_just = 0, axis_title_size = 12) + 
  scale_colour_manual(values = colors[c(2,4,6)]) +
  guides(fill=FALSE, color=FALSE)

#Fig2C: Number of Days to Sexual Metamorphosis
ggplot(expanded_T, aes(x=as.factor(Index), y=DPA, fill=as.factor(Index))) + 
  ylim(0,450) + 
  ylab("Time to maturation (days)") + 
  xlab("Initial Segment Count") + geom_violin(adjust=0.75, alpha=0.5) + 
  geom_boxplot(width=0.1, outlier.size=1, outlier.shape=21) + 
  stat_summary(fun="mean", geom="point", shape=23, size=5) + 
  scale_fill_manual(values=colors) + 
  geom_quasirandom(size=5, alpha=0.7, aes(color=Index)) + 
  scale_color_manual(values=colors) + 
  guides(fill=FALSE, color=FALSE) +
  theme_ipsum(axis_title_just = 0, axis_title_size = 12)
```
