## Supplementary File 02 for "The cost and payout of age on germline regeneration and sexual maturation in *Platynereis dumerilii*": Metzger_Fig3.R

```
#Figure 3:

#Loading packages (used across all figures)
library(ggplot2)
library(gcookbook)
library(dplyr)
library(hrbrthemes)
library(ggpubr)
library(ggsignif)
library(tidyr)
library(tidyverse)
library(reshape)
library(ggbeeswarm)
hrbrthemes::import_roboto_condensed()

#Setting color schemes
colors = c("#71254F","#a23671", "#B2793D", "#ffae58", "#317B7D", "#47B1B4", "#2F6A3A", "#93CF5B")

#Set working directory
setwd("~/Documents/Ozpolat/Experiments/Amputation & Fecundity")

#Reading in data
singles = read.csv("singles.csv", header=T, na.strings="")

##Making subsets
#subset data: T4
sing_T4 = subset(singles, (Timepoint %in% c("T4")))
#only survivors
sing_T4_surv = sing_T4[which(is.na(sing_T4$Days.surviving)),]
#only mortalities
sing_T4_mort = sing_T4[which(!is.na(sing_T4$Days.surviving)),]

#subset data: T3
sing_T3 = subset(singles, (Timepoint %in% c("T3")))
#survivors
sing_T3_surv = sing_T3[which(is.na(sing_T3$Days.surviving)),]
#only mortalities
sing_T3_mort = sing_T3[which(!is.na(sing_T3$Days.surviving)),]

#subset data: T2
sing_T2 = subset(singles, (Timepoint %in% c("T2")))
#survivors
sing_T2_surv = sing_T2[which(is.na(sing_T2$Days.surviving)),]
#only mortalities
sing_T2_mort = sing_T2[which(!is.na(sing_T2$Days.surviving)),]

#subset data: T1
sing_T1 = subset(singles, (Timepoint %in% c("T1")))
#survivors
sing_T1_surv = sing_T1[which(is.na(sing_T1$Days.surviving)),]
#only mortalities
sing_T1_mort = sing_T1[which(!is.na(sing_T1$Days.surviving)),]

#Fig3A
#T4: 40s control
ggplot(sing_T4[which(sing_T4$Index == "a"),], aes(x=as.numeric(DPA), y=as.numeric(Segments.added), group=Worm.ID, color=Worm.ID)) +
  #geom_point(shape=1) +
  geom_line(alpha=0.3, color=colors[1]) +
  stat_summary(aes(group = 1), geom = "point", fun.y = mean, shape = 15, size = 3, color=colors[1]) +ylim(0,60) +  
  stat_smooth(aes(group = 1), color=colors[1]) +
  xlab("DPA") +
  ylab("Segments Added") +
  theme_ipsum(axis_title_just = 0, axis_title_size =20, axis_text_size = 20) + 
  guides(fill=FALSE, color=FALSE)
length(unique(sing_T4[which(sing_T4$Index == "a"),]$Worm.ID)) #n=14

#T4: 60s control
ggplot(sing_T4[which(sing_T4$Index == "c"),], aes(x=as.numeric(DPA), y=as.numeric(Segments.added), group=Worm.ID, color=Worm.ID)) +
  #geom_point(shape=1) +
  geom_line(alpha=0.3, color=colors[5]) +
  stat_summary(aes(group = 1), geom = "point", fun.y = mean, shape = 17, size = 3, color=colors[5]) +ylim(0,60) +  
  stat_smooth(aes(group = 1), color=colors[5]) +
  xlab("DPA") +
  ylab("Segments Added") +
  theme_ipsum(axis_title_just = 0, axis_title_size =20, axis_text_size = 20) + 
  guides(fill=FALSE, color=FALSE)
length(unique(sing_T4[which(sing_T4$Index == "c"),]$Worm.ID)) #n=14

#Fig3B
#T4: 60s amputees
ggplot(sing_T4[which(sing_T4$Index == "d"),], aes(x=as.numeric(DPA), y=as.numeric(Segments.added), group=Worm.ID, color=Worm.ID)) +
  geom_line(alpha=0.3, color=colors[6]) +
  stat_summary(aes(group = 1), geom = "point", fun.y = mean, shape = 17, size = 3, color=colors[6]) +
  ylim(0,60) +  
  stat_smooth(aes(group = 1), color=colors[6]) +
  xlab("DPA") +
  ylab("Segments Added") +
  theme_ipsum(axis_title_just = 0, axis_title_size =20, axis_text_size = 20) + 
  guides(fill=FALSE, color=FALSE)
length(unique(sing_T4[which(sing_T4$Index == "d"),]$Worm.ID)) #n=38

#T4: 40s amputated
ggplot(sing_T4[which(sing_T4$Index == "b"),], aes(x=as.numeric(DPA), y=as.numeric(Segments.added), group=Worm.ID, color=Worm.ID)) +
  #geom_point(shape=1) +
  geom_line(alpha=0.3, color=colors[2]) +
  stat_summary(aes(group = 1), geom = "point", fun.y = mean, shape = 15, size = 3, color=colors[2]) +ylim(0,60) +  
  stat_smooth(aes(group = 1), color=colors[2]) +
  xlab("DPA") +
  ylab("Segments Added") +
  theme_ipsum(axis_title_just = 0, axis_title_size =20, axis_text_size = 20) + 
  guides(fill=FALSE, color=FALSE)
length(unique(sing_T4[which(sing_T4$Index == "b"),]$Worm.ID)) #n=15

#Fig3C
sing_7DPA = subset(singles, (DPA %in% c("7")))
#Graph
ggplot(sing_7DPA, aes(x=as.factor(Index), y=Segments.added)) +
  geom_quasirandom(size=5, alpha=0.7, aes(color=Index)) +
  geom_violin(adjust=1, alpha=0.5, aes(fill=Index)) +
  geom_boxplot(width=0.1, outline.size=1, outlier.shape=21, aes(fill=Index)) +
  stat_summary(fun="mean", geom="point", shape=23, size=3, alpha=0.3, aes(fill=Index)) +
  theme_ipsum(axis_title_just = 0, axis_title_size =20, axis_text_size = 20) + 
  scale_fill_manual(values=colors[c(1,2,5,6)]) + 
  scale_color_manual(values=colors[c(1,2,5,6)]) + 
  xlab("Group") +
  ylab("Segments Added") +
  ylim(-7,55) +
  guides(fill=FALSE, color=FALSE)
#ANOVA - factorial
dpa7_aov = aov(Segments.added ~ Segment.group * Treatment, data = sing_7DPA)
summary(dpa7_aov)
#yes, significant: Segment.group significant, Treatment not, interaction not
pairwise.t.test(sing_7DPA$Segments.added, sing_7DPA$Index, p.adjust.method = "BH")
#n's
length(which(sing_7DPA$Index=="a")) #46
length(which(sing_7DPA$Index=="b")) #49
length(which(sing_7DPA$Index=="c")) #44
length(which(sing_7DPA$Index=="d")) #71

#Fig3D
sing_14DPA = subset(singles, (DPA %in% c("14")))
#Graph
ggplot(sing_14DPA, aes(x=as.factor(Index), y=Segments.added)) +
  geom_quasirandom(size=5, alpha=0.7, aes(color=Index)) +
  geom_violin(adjust=1, alpha=0.5, aes(fill=Index)) +
  geom_boxplot(width=0.1, outline.size=1, outlier.shape=21, aes(fill=Index)) +
  stat_summary(fun="mean", geom="point", shape=23, size=5, aes(fill=Index)) +
  theme_ipsum(axis_title_just = 0, axis_title_size =20, axis_text_size = 20) + 
  scale_fill_manual(values=colors[c(1,2,5,6)]) + 
  scale_color_manual(values=colors[c(1,2,5,6)]) + 
  xlab("Group") +
  ylab("Segments Added") +
  ylim(-7,55) +
  guides(fill=FALSE, color=FALSE)
#ANOVA - factorial
dpa14_aov = aov(Segments.added ~ Segment.group * Treatment, data = sing_14DPA)
summary(dpa14_aov)
#yes, significant: Segment.group not significant, Treatment significant, interaction not
pairwise.t.test(sing_14DPA$Segments.added, sing_14DPA$Index, p.adjust.method = "BH")
#n's
length(which(sing_14DPA$Index=="a")) #19
length(which(sing_14DPA$Index=="b")) #10
length(which(sing_14DPA$Index=="c")) #20
length(which(sing_14DPA$Index=="d")) #38

#Fig3E
sing_28DPA = subset(singles, (DPA %in% c("28","29","30","31","32")))
#Graph
ggplot(sing_28DPA, aes(x=as.factor(Index), y=Segments.added)) +
  geom_quasirandom(size=5, alpha=0.7, aes(color=Index)) +
  geom_violin(adjust=1, alpha=0.5, aes(fill=Index)) +
  geom_boxplot(width=0.1, outline.size=1, outlier.shape=21, aes(fill=Index)) +
  stat_summary(fun="mean", geom="point", shape=23, size=5, aes(fill=Index)) +
  theme_ipsum(axis_title_just = 0, axis_title_size =20, axis_text_size = 20) + 
  scale_fill_manual(values=colors[c(1,2,5,6)]) + 
  scale_color_manual(values=colors[c(1,2,5,6)]) + 
  xlab("Group") +
  ylab("Segments Added") +
  ylim(-7,55) +
  guides(fill=FALSE, color=FALSE)
#ANOVA - factorial
dpa28_aov = aov(Segments.added ~ Segment.group * Treatment, data = sing_28DPA)
summary(dpa28_aov)
#yes, significant: Segment.group not significant, Treatment significant, interaction significant
pairwise.t.test(sing_28DPA$Segments.added, sing_28DPA$Index, p.adjust.method = "BH")
#n's
length(which(sing_28DPA$Index=="a")) #29
length(which(sing_28DPA$Index=="b")) #30
length(which(sing_28DPA$Index=="c")) #30
length(which(sing_28DPA$Index=="d")) #32

#Fig3F
sing_63DPA = subset(singles, (DPA %in% c("63","64")))
#Graph
ggplot(sing_63DPA, aes(x=as.factor(Index), y=Segments.added)) +
  geom_quasirandom(size=5, alpha=0.7, aes(color=Index)) +
  geom_violin(adjust=1, alpha=0.5, aes(fill=Index)) +
  geom_boxplot(width=0.1, outline.size=1, outlier.shape=21, aes(fill=Index)) +
  stat_summary(fun="mean", geom="point", shape=23, size=5, aes(fill=Index)) +
  theme_ipsum(axis_title_just = 0, axis_title_size =20, axis_text_size = 20) + 
  scale_fill_manual(values=colors[c(1,2,5,6)]) + 
  scale_color_manual(values=colors[c(1,2,5,6)]) + 
  xlab("Group") +
  ylab("Segments Added") +
  ylim(-7,55) +
  guides(fill=FALSE, color=FALSE)
#ANOVA - factorial
dpa63_aov = aov(Segments.added ~ Segment.group * Treatment, data = sing_63DPA)
summary(dpa63_aov)
#yes, significant: Segment.group not significant, Treatment significant, interaction significant
pairwise.t.test(sing_63DPA$Segments.added, sing_63DPA$Index, p.adjust.method = "BH")
#n's
length(which(sing_63DPA$Index=="a")) #14
length(which(sing_63DPA$Index=="b")) #14
length(which(sing_63DPA$Index=="c")) #12
length(which(sing_63DPA$Index=="d")) #14
```
