## Supplementary File 02 for "The cost and payout of age on germline regeneration and sexual maturation in *Platynereis dumerilii*": Metzger_Fig4.R

```
#Figure 4:

#Loading packages (used across all figures)
library(ggplot2)
library(gcookbook)
library(dplyr)
library(hrbrthemes)
library(ggpubr)
library(ggsignif)
library(tidyr)
library(tidyverse)
library(reshape)
library(ggbeeswarm)
hrbrthemes::import_roboto_condensed()

#Setting color schemes
colors = c("#71254F","#a23671", "#B2793D", "#ffae58", "#317B7D", "#47B1B4", "#2F6A3A", "#93CF5B")

#Set working directory
setwd("~/Documents/Ozpolat/Experiments/Amputation & Fecundity")

#Reading in data
singles = read.csv("singles.csv", header=T, na.strings="")

##Making subsets
#subset data: T4
sing_T4 = subset(singles, (Timepoint %in% c("T4")))
#only survivors
sing_T4_surv = sing_T4[which(is.na(sing_T4$Days.surviving)),]
#only mortalities
sing_T4_mort = sing_T4[which(!is.na(sing_T4$Days.surviving)),]
just_morts = sing_T4_mort %>% distinct(Worm.ID, .keep_all = T)

#Fig4A
#PROPORTIONS OF DEATH
piechart = unique(sing_T4[,c(3,8,20)]) #only T4
#40s ctrls
total_40_ctrl = length(which(piechart$Index=="a")) #14
dead_40_ctrl = length(which(piechart$Index=="a" & !is.na(piechart$Days.surviving)))
x = c((dead_40_ctrl/total_40_ctrl)*100, 100-((dead_40_ctrl/total_40_ctrl)*100))
labels = c("Died","Survived")
pie(x,labels)
dead_40_ctrl/total_40_ctrl*100 #0

#40s amps
total_40_amp = length(which(piechart$Index=="b")) #16
dead_40_amp = length(which(piechart$Index=="b" & !is.na(piechart$Days.surviving)))
x = c((dead_40_amp/total_40_amp)*100, 100-((dead_40_amp/total_40_amp)*100))
labels = c("Died","Survived")
pie(x,labels)
dead_40_amp/total_40_amp*100 #6.25

#60s ctrls
total_60_ctrl = length(which(piechart$Index=="c")) #15
dead_60_ctrl = length(which(piechart$Index=="c" & !is.na(piechart$Days.surviving)))
x = c((dead_60_ctrl/total_60_ctrl)*100, 100-((dead_60_ctrl/total_60_ctrl)*100))
labels = c("Died","Survived")
pie(x,labels)
dead_60_ctrl/total_60_ctrl*100 #20

#60s amps
total_60_amp = length(which(piechart$Index=="d")) #40
dead_60_amp = length(which(piechart$Index=="d" & !is.na(piechart$Days.surviving)))
x = c((dead_60_amp/total_60_amp)*100, 100-((dead_60_amp/total_60_amp)*100))
labels = c("Died","Survived")
pie(x,labels)
dead_60_amp/total_60_amp*100 #57.5

#Fig4B
ggplot(just_morts, aes(x = Days.surviving, fill = Index)) +
  scale_fill_manual(values=colors[c(2,5,6)]) +
  geom_histogram(position = "stack", alpha = 1) +
  xlab("Days Surviving") +
  ylab("Count") +
  theme_ipsum(axis_title_just = 0, axis_title_size =20, axis_text_size = 20) +
  guides(fill=FALSE)

#Fig4C
#get a list of unique dead IDs
mort_IDS = just_morts$Worm.ID
#for worm in list, get a list of segments added, return maximum, add to vector
seg_add_vector = c()
for (worm in mort_IDS) {
  ID_indices = which(sing_T4_mort$Worm.ID==worm) #list of indices for ID
  max_added = max(sing_T4_mort$Segments.added[c(ID_indices)])
  seg_add_vector = c(seg_add_vector,max_added)
}
seg_add_vector[is.na(seg_add_vector)] = 0 #transform NA to 0 (NA means they did not get past posterior regeneration)
#transform df
just_morts$Segments.added = seg_add_vector
#plot
ggplot(just_morts, aes(Segments.added, Days.surviving, color=Index, fill=Index, alpha=0.5,size=10, shape=Index)) +
  geom_point() +
  theme_ipsum() + 
  scale_color_manual(values=colors[c(2,5,6)])+
  scale_fill_manual(values=colors[c(2,5,6)]) +
  xlab("") +
  ylab("") +
  guides(fill=FALSE, color=FALSE, alpha=FALSE, size=FALSE, shape=FALSE)

#Fig4D
#original segment number & days surviving
ggplot(just_morts, aes(Original.segments, Days.surviving, color=Index, fill=Index, alpha=0.5,size=10, shape=Index)) +
  geom_point() +
  xlim(60, 70) +
  theme_ipsum() + 
  scale_color_manual(values=colors[c(2,5,6)])+ #bcd
  scale_fill_manual(values=colors[c(2,5,6)]) +
  #xlab("") +
  #ylab("") +
  guides(fill=FALSE, color=FALSE, alpha=FALSE, size=FALSE, shape=FALSE)
#note: error msg is throwing out one 40s amp point (did not subset, limited axes)
```
