## Supplementary File 02 for "The cost and payout of age on germline regeneration and sexual maturation in *Platynereis dumerilii*": Metzger_Fig5.R

```
#Figure 5:

#Loading packages (used across all figures)
library(ggplot2)
library(gcookbook)
library(dplyr)
library(hrbrthemes)
library(ggpubr)
library(ggsignif)
library(tidyr)
library(tidyverse)
library(reshape)
library(ggbeeswarm)
hrbrthemes::import_roboto_condensed()

#Setting color schemes
colors = c("#71254F","#a23671", "#B2793D", "#ffae58", "#317B7D", "#47B1B4", "#2F6A3A", "#93CF5B")

#Set working directory
setwd("~/Documents/Ozpolat/Experiments/Amputation & Fecundity")

#Reading in data
singles = read.csv("singles.csv", header=T, na.strings="")

#regeneration staging: proportional bar charts
#NOTE: recolored in Adobe illustrator
regens = singles[complete.cases(singles$Stage.of.blastema.letter),]
#3DPA
reg_3DPA = subset(regens, (DPA %in% c(3)))
ggplot(reg_3DPA, aes(x = Index, fill = Stage.of.blastema.letter)) +
  geom_bar(position = "fill", color="black") +
  scale_y_continuous(labels = scales::percent) +
  scale_fill_manual(values=greenscale[c(1,2,3)]) +
  theme_ipsum(axis_title_just = 0, axis_title_size =20, axis_text_size = 20) +
  guides(fill=FALSE)
length(which(reg_3DPA$Index=="b")) #9
length(which(reg_3DPA$Index=="d")) #14

#7DPA
reg_7DPA = subset(regens, (DPA %in% c(7)))
ggplot(reg_7DPA, aes(x = Index, fill = Stage.of.blastema.letter)) +
  geom_bar(position = "fill", color="black") +
  scale_y_continuous(labels = scales::percent) +
  scale_fill_manual(values=greenscale[c(1,2,4,5,6,7)]) +
  theme_ipsum(axis_title_just = 0, axis_title_size =20, axis_text_size = 20) +
  guides(fill=FALSE)
length(which(reg_7DPA$Index=="b")) #47
length(which(reg_7DPA$Index=="d")) #31

#14DPA
reg_14DPA = subset(regens, (DPA %in% c(14)))
ggplot(reg_14DPA, aes(x = Index, fill = Stage.of.blastema.letter)) +
  geom_bar(position = "fill", color="black") +
  scale_y_continuous(labels = scales::percent) +
  scale_fill_manual(values=greenscale[c(2,6,1,7)]) +
  theme_ipsum(axis_title_just = 0, axis_title_size =20, axis_text_size = 20) +
  guides(fill=FALSE)
length(which(reg_14DPA$Index=="b")) #10
length(which(reg_14DPA$Index=="d")) #37
```
