## Supplementary File 02 for "The cost and payout of age on germline regeneration and sexual maturation in *Platynereis dumerilii*": Metzger_FigS1.R

```
##Supplemental Figure 1:

#Loading packages
library(ggplot2)
library(gcookbook)
library(dplyr)
library(hrbrthemes)
library(ggpubr)
library(ggsignif)
library(tidyr)
library(tidyverse)
library(reshape)
library(ggbeeswarm)
hrbrthemes::import_roboto_condensed()

#Setting color schemes
colors = c("#71254F","#a23671", "#B2793D", "#ffae58", "#317B7D", "#47B1B4", "#2F6A3A", "#93CF5B")
purples = c("#3F1E5F","#6A329F","#B498CF")

#Defining functions
theme_blank <- function(...) {
  ret <- theme_bw(...)
  ret$line <- element_blank()
  ret$rect <- element_blank()
  ret$strip.text <- element_blank()
  ret$axis.text <- element_blank()
  ret$plot.title <- element_blank()
  ret$axis.title <- element_blank()
  ret$plot.margin <- structure(c(0, 0, -1, -1), unit = "lines", valid.unit = 3L, class = "unit")
  ret
}
ggplotRegression <- function (fit, title, clr, x_lab, y_lab) {
  
  require(ggplot2)
  
  ggplot(fit$model, aes_string(x = names(fit$model)[2], y = names(fit$model)[1])) + 
    geom_point() +
    theme_ipsum(axis_title_just = 0, axis_title_size =20, axis_text_size = 20) +
    xlab(x_lab) +
    ylab(y_lab) +
    stat_smooth(method = "lm", col = clr) +
    labs(title = paste("Adj R2 = ",signif(summary(fit)$adj.r.squared, 5),
                       "Intercept =",signif(fit$coef[[1]],5 ),
                       " Slope =",signif(fit$coef[[2]], 5),
                       " P =",signif(summary(fit)$coef[2,4], 5)))
}

#Set wd
setwd("~/Documents/Ozpolat/Experiments/Amputation & Fecundity/")

#reading in data
df1 = read.csv(file="amp_pilot.csv", header=TRUE)
AE = read.csv(file="AE.csv", header=T, sep=",")


#FigS1A: 1 DPA
ggplot(df1[which(df1$DPA==1),], aes(x=as.factor(Index), y=Stage)) +
  geom_quasirandom(size=5, alpha=0.7, aes(color=Index)) +
  geom_violin(adjust=1, alpha=0.5, aes(fill=Index)) +
  geom_boxplot(width=0.1, outline.size=1, outlier.shape=21, aes(fill=Index)) +
  stat_summary(fun="mean", geom="point", shape=23, size=5, aes(fill=Index)) +
  theme_ipsum(axis_title_just = 0, axis_title_size =20, axis_text_size = 20) + 
  scale_fill_manual(values=purples[c(1,2,3)]) + 
  scale_color_manual(values=purples[c(1,2,3)]) + 
  xlab("Group") +
  ylab("Segments Added") +
  ylim(0,6) +
  guides(fill=FALSE, color=FALSE)
#anova
dpa1 = df1[which(df1$DPA==1),]
dpa1_anova = aov(Stage ~ Index, data = dpa1) #univariate anova
summary(dpa1_anova) #different!
TukeyHSD(dpa1_anova, conf.level=0.95)
#not significant: b-a
#significant: c-a, c-b

#FigS1A: 3 DPA
ggplot(df1[which(df1$DPA==3),], aes(x=as.factor(Index), y=Stage)) +
  geom_quasirandom(size=5, alpha=0.7, aes(color=Index)) +
  geom_violin(adjust=1, alpha=0.5, aes(fill=Index)) +
  geom_boxplot(width=0.1, outline.size=1, outlier.shape=21, aes(fill=Index)) +
  stat_summary(fun="mean", geom="point", shape=23, size=5, aes(fill=Index)) +
  theme_ipsum(axis_title_just = 0, axis_title_size =20, axis_text_size = 20) + 
  scale_fill_manual(values=purples[c(1,2,3)]) + 
  scale_color_manual(values=purples[c(1,2,3)]) + 
  xlab("Group") +
  ylab("Segments Added") +
  ylim(0,6) +
  guides(fill=FALSE, color=FALSE)
#anova
dpa3 = df1[which(df1$DPA==3),]
dpa3_anova = aov(Stage ~ Index, data = dpa3) #univariate anova
summary(dpa3_anova) #different!
TukeyHSD(dpa3_anova, conf.level=0.95)
#not significant: none
#significant: c-a, c-b, b-a

#FigS1A: 5 DPA
ggplot(df1[which(df1$DPA==5),], aes(x=as.factor(Index), y=Stage)) +
  geom_quasirandom(size=5, alpha=0.7, aes(color=Index)) +
  geom_violin(adjust=1, alpha=0.5, aes(fill=Index)) +
  geom_boxplot(width=0.1, outline.size=1, outlier.shape=21, aes(fill=Index)) +
  stat_summary(fun="mean", geom="point", shape=23, size=5, aes(fill=Index)) +
  theme_ipsum(axis_title_just = 0, axis_title_size =20, axis_text_size = 20) + 
  scale_fill_manual(values=purples[c(1,2,3)]) + 
  scale_color_manual(values=purples[c(1,2,3)]) + 
  xlab("Group") +
  ylab("Segments Added") +
  ylim(0,6) +
  guides(fill=FALSE, color=FALSE)
#anova
dpa5 = df1[which(df1$DPA==5),]
dpa5_anova = aov(Stage ~ Index, data = dpa5) #univariate anova
summary(dpa5_anova) #different!
TukeyHSD(dpa5_anova, conf.level=0.95)
#not significant: b-a
#significant: c-a, c-b

#FigS1A: 7 DPA
ggplot(df1[which(df1$DPA==7),], aes(x=as.factor(Index), y=Stage)) +
  geom_quasirandom(size=5, alpha=0.7, aes(color=Index)) +
  geom_violin(adjust=1, alpha=0.5, aes(fill=Index)) +
  geom_boxplot(width=0.1, outline.size=1, outlier.shape=21, aes(fill=Index)) +
  stat_summary(fun="mean", geom="point", shape=23, size=5, aes(fill=Index)) +
  theme_ipsum(axis_title_just = 0, axis_title_size =20, axis_text_size = 20) + 
  scale_fill_manual(values=purples[c(1,2,3)]) + 
  scale_color_manual(values=purples[c(1,2,3)]) + 
  xlab("Group") +
  ylab("Segments Added") +
  ylim(0,6) +
  guides(fill=FALSE, color=FALSE)
#anova
dpa7 = df1[which(df1$DPA==7),]
dpa7_anova = aov(Stage ~ Index, data = dpa7) #univariate anova
summary(dpa7_anova) #different!
TukeyHSD(dpa7_anova, conf.level=0.95)
#not significant: b-a, c-b
#significant: c-a

#FigS1B
AE_male = subset(AE, (Sex %in% c("Male")))
ggplot(AE_male, aes(x=as.factor(Index), y=Segments)) +
  geom_quasirandom(size=5, alpha=0.7, aes(color=Index)) +
  geom_violin(adjust=1, alpha=0.5, aes(fill=Index)) +
  geom_boxplot(width=0.1, outlier.shape=21, aes(fill=Index)) +
  stat_summary(fun="mean", geom="point", shape=23, size=5, aes(fill=Index)) +
  theme_ipsum(axis_title_just = 0, axis_title_size =20, axis_text_size = 20) + 
  scale_fill_manual(values=colors) + 
  scale_color_manual(values=colors) + 
  xlab("Group") +
  ylab("Segments") +
  ylim(60,85) +
  guides(fill=FALSE, color=FALSE)

#FigS1C
AE_female = subset(AE, (Sex %in% c("Female")))
ggplot(AE_female, aes(x=as.factor(Index), y=Segments)) +
  geom_quasirandom(size=5, alpha=0.7, aes(color=Index)) +
  geom_violin(adjust=1, alpha=0.5, aes(fill=Index)) +
  geom_boxplot(width=0.1, outlier.shape=21, aes(fill=Index)) +
  stat_summary(fun="mean", geom="point", shape=23, size=5, aes(fill=Index)) +
  theme_ipsum(axis_title_just = 0, axis_title_size =20, axis_text_size = 20) + 
  scale_fill_manual(values=colors) + 
  scale_color_manual(values=colors) + 
  xlab("Group") +
  ylab("Segments") +
  ylim(60,85) +
  guides(fill=FALSE, color=FALSE)

#FigS1D
#AE without NA values
AE_noNA = AE[complete.cases(AE),]
x = AE_noNA$Days.since.start
y = AE_noNA$Segments
lmat_model = lm(y ~ x)
summary(lmat_model)
ggplot(AE_noNA, aes(x, y)) +
  geom_point(size = 5, alpha = 0.9, aes(color=Group)) +
  scale_color_manual(values=colors) +
  geom_smooth(method='lm', color="black") +
  labs(title = paste("Adj R2 = ",signif(summary(lmat_model)$adj.r.squared, 5),
                     "Intercept =",signif(lmat_model$coef[[1]],5 ),
                     " Slope =",signif(lmat_model$coef[[2]], 5),
                     " P =",signif(summary(lmat_model)$coef[2,4], 5))) +
  theme_ipsum(axis_title_just = 0, axis_title_size =20, axis_text_size = 20) + 
  guides(color=FALSE)
```
