## Supplementary File 02 for "The cost and payout of age on germline regeneration and sexual maturation in *Platynereis dumerilii*": Metzger_FigS2.R

```
#Figure 2S:

#Loading packages (used across all figures)
library(ggplot2)
library(gcookbook)
library(dplyr)
library(hrbrthemes)
library(ggpubr)
library(ggsignif)
library(tidyr)
library(tidyverse)
library(reshape)
library(ggbeeswarm)
hrbrthemes::import_roboto_condensed()

#Setting color schemes
colors = c("#71254F","#a23671", "#B2793D", "#ffae58", "#317B7D", "#47B1B4", "#2F6A3A", "#93CF5B")
cscheme = c("#a23671", "#71254F", "#ffae58", "#B2793D", "#47B1B4", "#317B7D")

#Set working directory
setwd("~/Documents/Ozpolat/Experiments/Amputation & Fecundity")

#Reading in data
format = read.csv("forMAT-final.csv", header=T) #cumulative maturations, not separated by box
death_cumulative = read.csv(file="death_cumulative.csv", header = TRUE, sep = ",")
wk4 = read.csv("AF_4wk_scounts.csv")
wk8 = read.csv("AF_8wk_scounts.csv")
wk12 = read.csv("AF_12wk_scounts.csv")

#Subsets
format_60s = format[which(format$Treatment=="s60_amputated"|format$Treatment=="s60_control"),]
format_50s = format[which(format$Treatment=="s50_amputated"|format$Treatment=="s50_control"),]
format_40s = format[which(format$Treatment=="s40_amputated"|format$Treatment=="s40_control"),]
#FigS2A
ggplot(format_40s, aes(x=DPA, y = Percent_M, fill = as.factor(Index), colour = Index)) + 
  ylim(0, 60) + 
  xlim(0,360) + 
  xlab("Days Post Amputation (DPA)") + 
  ylab("Percent Mature") + 
  geom_line(size=1.5) +
  theme_ipsum(axis_title_just = 0, axis_title_size = 12) + 
  scale_colour_manual(values = c(colors[1], colors[2])) + 
  guides(color=FALSE)
#FigS2A'
ggplot(format_50s, aes(x=DPA, y = Percent_M, fill = as.factor(Index), colour = Index)) + 
  ylim(0, 60) + 
  xlim(0,360) + 
  xlab("Days Post Amputation (DPA)") + 
  ylab("Percent Mature") + 
  geom_line(size=1.5) +
  theme_ipsum(axis_title_just = 0, axis_title_size = 12) + 
  scale_colour_manual(values = c(colors[3], colors[4])) + 
  guides(color=FALSE)
#FigS2A"
ggplot(format_60s, aes(x=DPA, y = Percent_M, fill = as.factor(Index), colour = Index)) + 
  ylim(0, 60) + 
  xlim(0,360) + 
  xlab("Days Post Amputation (DPA)") + 
  ylab("Percent Mature") + 
  geom_line(size=1.5) +
  theme_ipsum(axis_title_just = 0, axis_title_size = 12) + 
  scale_colour_manual(values = c(colors[5], colors[6])) + 
  guides(color=FALSE)

#Making subsets
deathc_60s = death_cumulative[which(death_cumulative$Treatment=="s60"),]
deathc_50s = death_cumulative[which(death_cumulative$Treatment=="s50"),]
deathc_40s = death_cumulative[which(death_cumulative$Treatment=="s40"),]

#FigS2B
ggplot(deathc_40s, aes(x = days, y = ((Perc)*-1), fill = as.factor(Condition), colour=Condition)) + 
  ylim(-100, 0) + 
  xlab("Days Post Amputation (DPA)") + 
  ylab("Percent surviving") + 
  geom_line(size=1.5) +
  theme_ipsum(axis_title_just = 0, axis_title_size = 12) + 
  scale_colour_manual(values=c(colors[2],colors[1])) + 
  guides(color=F)
#FigS2B'
ggplot(deathc_50s, aes(x = days, y = ((Perc)*-1), fill = as.factor(Condition), colour=Condition)) + 
  ylim(-100, 0) + 
  xlab("Days Post Amputation (DPA)") + 
  ylab("Percent surviving") + 
  geom_line(size=1.5) +
  theme_ipsum(axis_title_just = 0, axis_title_size = 12) + 
  scale_colour_manual(values=c(colors[4],colors[3])) + 
  guides(color=F)
#FigS2B"
ggplot(deathc_60s, aes(x = days, y = ((Perc)*-1), fill = as.factor(Condition), colour=Condition)) + 
  ylim(-100, 0) + 
  xlab("Days Post Amputation (DPA)") + 
  ylab("Percent surviving") + 
  geom_line(size=1.5) +
  theme_ipsum(axis_title_just = 0, axis_title_size = 12) + 
  scale_colour_manual(values=c(colors[6],colors[5])) + 
  guides(color=F)

#FigS2C
#4wk
#n's
length(which(wk4$treatment=="s40_amputated")) #62
length(which(wk4$treatment=="s50_amputated")) #58
length(which(wk4$treatment=="s60_amputated")) #35
ggplot(wk4, aes(x=as.factor(treatment), y=counts)) + 
  ylim(0,90) + 
  xlab("Initial Segment Count") + 
  ylab("28 DPA Segment Count") + 
  geom_quasirandom(size=5, alpha=0.5, aes(color=treatment)) +
  geom_violin(adjust=0.5, alpha=0.5, aes(fill=treatment)) + 
  geom_boxplot(width=0.1, outlier.size=0, outlier.shape=21, aes(fill=treatment)) + 
  stat_summary(fun="mean", geom="point", shape=23, size=5, aes(fill=treatment)) + 
  theme_ipsum(axis_title_just = 0, axis_title_size =12) + 
  scale_fill_manual(values=c("#a23671", "#ffae58", "#47b1b4")) + 
  scale_color_manual(values=c("#a23671", "#ffae58", "#47b1b4")) + 
  guides(fill=FALSE, color=FALSE)
#FigS2C'
#8wk
#n's
length(which(wk8$treatment=="s40_control")) #29
length(which(wk8$treatment=="s40_amputated")) #49
length(which(wk8$treatment=="s50_control")) #40
length(which(wk8$treatment=="s50_amputated")) #46
length(which(wk8$treatment=="s60_control")) #18
length(which(wk8$treatment=="s60_amputated")) #21
ggplot(wk8, aes(x=as.factor(treatment), y=counts)) + 
  ylim(0,90) + 
  xlab("Initial Segment Count") + 
  ylab("Segment Count") + 
  geom_quasirandom(size=5, alpha=0.5, aes(color=treatment)) +
  geom_violin(adjust=0.5, alpha=0.5, aes(fill=treatment)) + 
  geom_boxplot(width=0.1, outlier.size=0, outlier.shape=21, aes(fill=treatment)) + 
  stat_summary(fun="mean", geom="point", shape=23, size=5, aes(fill=treatment)) +
  theme_ipsum(axis_title_just = 0, axis_title_size =12) + 
  scale_fill_manual(values=cscheme) + 
  scale_color_manual(values=cscheme) + 
  guides(fill=FALSE, color=FALSE)
#FigS2C"
#12wk
#n's
length(which(wk12$treatment=="s40_control")) #44
length(which(wk12$treatment=="s40_amputated")) #47
length(which(wk12$treatment=="s50_control")) #35
length(which(wk12$treatment=="s50_amputated")) #38
length(which(wk12$treatment=="s60_control")) #18
length(which(wk12$treatment=="s60_amputated")) #20
ggplot(wk12, aes(x=as.factor(treatment), y=counts)) + 
  ylim(0,90) + 
  xlab("Initial Segment Count") + 
  ylab("Segment Count") + 
  geom_quasirandom(size=5, alpha=0.5, aes(color=treatment)) +
  geom_violin(adjust=0.5, alpha=0.5, aes(fill=treatment)) + 
  geom_boxplot(width=0.1, outlier.size=0, outlier.shape=21, aes(fill=treatment)) + 
  stat_summary(fun="mean", geom="point", shape=23, size=5, aes(fill=treatment)) +
  theme_ipsum(axis_title_just = 0, axis_title_size =12) + 
  scale_fill_manual(values=cscheme) + 
  scale_color_manual(values=cscheme) + 
  guides(fill=FALSE, color=FALSE)
#statistical tests
#wk4
#two tailed t test between 40S and 50S
t.test(x=wk4$counts[which(wk4$treatment=="s40_amputated")], y =wk4$counts[which(wk4$treatment=="s50_amputated")], alternative = "two.sided", var.equal = FALSE) #pval = 0.0004267 ***
#two tailed t test between 40S and 60S
t.test(x=wk4$counts[which(wk4$treatment=="s40_amputated")], y =wk4$counts[which(wk4$treatment=="s60_amputated")], alternative = "two.sided", var.equal = FALSE) #pval = 0.003771 **
#two tailed t test between 50S and 60S
t.test(x=wk4$counts[which(wk4$treatment=="s50_amputated")], y =wk4$counts[which(wk4$treatment=="s60_amputated")], alternative = "two.sided", var.equal = FALSE) #pval = 0.5696 NS

#wk8
#two tailed t test between amp and amp
t.test(x=wk8$counts[which(wk8$treatment=="s40_amputated")], y =wk8$counts[which(wk8$treatment=="s50_amputated")], alternative = "two.sided", var.equal = FALSE) #pval = 0.0005853 ***
#two tailed t test between 40S and 60S amp
t.test(x=wk8$counts[which(wk8$treatment=="s40_amputated")], y =wk8$counts[which(wk8$treatment=="s60_amputated")], alternative = "two.sided", var.equal = FALSE) #pval = 0.0001145 ***
#two tailed t test between 50S and 60S amp
t.test(x=wk8$counts[which(wk8$treatment=="s50_amputated")], y =wk8$counts[which(wk8$treatment=="s60_amputated")], alternative = "two.sided", var.equal = FALSE) #pval = 0.02905 *

#between controls
t.test(x=wk8$counts[which(wk8$treatment=="s40_amputated")], y =wk8$counts[which(wk8$treatment=="s40_control")], alternative = "two.sided", var.equal = FALSE) #pval = 5.194e-07 ***
#two tailed t test between 40S and 60S
t.test(x=wk8$counts[which(wk8$treatment=="s50_amputated")], y =wk8$counts[which(wk8$treatment=="s50_control")], alternative = "two.sided", var.equal = FALSE) #pval = 8.892e-08 ***
#two tailed t test between 50S and 60S
t.test(x=wk8$counts[which(wk8$treatment=="s60_amputated")], y =wk8$counts[which(wk8$treatment=="s60_control")], alternative = "two.sided", var.equal = FALSE) #pval = 0.1571 NS

#wk12
#two tailed t test between amp and amp
t.test(x=wk12$counts[which(wk12$treatment=="s40_amputated")], y =wk12$counts[which(wk12$treatment=="s50_amputated")], alternative = "two.sided", var.equal = FALSE) #pval = 0.02757 *
#two tailed t test between 40S and 60S amp
t.test(x=wk12$counts[which(wk12$treatment=="s40_amputated")], y =wk12$counts[which(wk12$treatment=="s60_amputated")], alternative = "two.sided", var.equal = FALSE) #pval = 0.0001229 ***
#two tailed t test between 50S and 60S amp
t.test(x=wk12$counts[which(wk12$treatment=="s50_amputated")], y =wk12$counts[which(wk12$treatment=="s60_amputated")], alternative = "two.sided", var.equal = FALSE) #pval = 0.05722 NS

#between controls
#two tailed t test between 40s
t.test(x=wk12$counts[which(wk12$treatment=="s40_amputated")], y =wk12$counts[which(wk12$treatment=="s40_control")], alternative = "two.sided", var.equal = FALSE) #pval = 5.976e-07 ***
#two tailed t test between 50s
t.test(x=wk12$counts[which(wk12$treatment=="s50_amputated")], y =wk12$counts[which(wk12$treatment=="s50_control")], alternative = "two.sided", var.equal = FALSE) #pval = 0.06031 NS
#two tailed t test between 60s
t.test(x=wk12$counts[which(wk12$treatment=="s60_amputated")], y =wk12$counts[which(wk12$treatment=="s60_control")], alternative = "two.sided", var.equal = FALSE) #pval = 0.1909 NS
```
