## Supplementary File 02 for "The cost and payout of age on germline regeneration and sexual maturation in *Platynereis dumerilii*": Metzger_FigS3.R

```
#Figure S3:

#Loading packages (used across all figures)
library(ggplot2)
library(gcookbook)
library(dplyr)
library(hrbrthemes)
library(ggpubr)
library(ggsignif)
library(tidyr)
library(tidyverse)
library(reshape)
library(ggbeeswarm)
hrbrthemes::import_roboto_condensed()

#Setting color schemes
colors = c("#71254F","#a23671", "#B2793D", "#ffae58", "#317B7D", "#47B1B4", "#2F6A3A", "#93CF5B")

#Set working directory
setwd("~/Documents/Ozpolat/Experiments/Amputation & Fecundity")

#Reading in data
t4_rates = read.csv("t4_rates.csv", header=T, na.strings="")

#FigS3A
t4_rates_alive = t4_rates[is.na(t4_rates$Days.surviving),]
ggplot(t4_rates_alive, aes(x=DPA, y=Rate, color=Index, fill=Index, shape=Index)) +
  geom_point(size=2, shape=20, aes(alpha=0.01)) +
  geom_smooth() +
  scale_color_manual(values=colors[c(1,2,5,6)])+
  scale_fill_manual(values=colors[c(1,2,5,6)]) +
  theme_ipsum() +
  xlab("DPA") +
  ylab("Rate of Segment Addition") +
  guides(fill=FALSE, color=FALSE, alpha=FALSE)

#n's
length(unique(subset(t4_rates_alive, Index == "a")$Worm.ID)) #14
length(unique(subset(t4_rates_alive, Index == "b")$Worm.ID)) #14
length(unique(subset(t4_rates_alive, Index == "c")$Worm.ID)) #12
length(unique(subset(t4_rates_alive, Index == "d")$Worm.ID)) #14
```
